# Female Asian elephant clans have age-based order but weak dominance resolution

**DOI:** 10.1101/2025.08.22.671838

**Authors:** S. Nandini, Hansraj Gautam, P. Keerthipriya, T.N.C. Vidya

**Affiliations:** Evolutionary and Organismal Biology Unit, Jawaharlal Nehru Centre for Advanced Scientific Research (JNCASR), Bengaluru, India, 560064

**Keywords:** Animal groups, socioecology, within-group competition, fission-fusion, agonistic interactions, dominance, association index, age, matriarchs, centrality

## Abstract

The variation in dominance relationships in group-living species is often interpreted through socioecological frameworks that link social structures to resource-risk distributions. However, in elephants, such inferences are hindered by a lack of comparable assessments of within-group dominance in different species. To advance our understanding of elephant socioecology, we present here, the first study on agonistic and dominance relationships within female Asian elephant clans (most inclusive social groups), and compare our results with those from African savannah elephants. By analysing agonistic interactions, and dominance and association networks based on over four years of observations of five clans, we show that Asian elephants have a resolved but weakly structured within-clan dominance order. Female dyads showed unidirectionality, but triad motif structures of dominance networks suggested resolved dominance only in some clans. Older females were more dominant although there were moderate levels of reversals against age-based order and age difference did not dampen dyadic conflict. Neither older age nor dominant status conferred more central status in the female association network. Weak dominance resolution and the effects of age contrast with the stronger dominance and age-based order found in African savannah elephants. We identify potential socioecological and demographic explanations of female dominance in elephants.

## Introduction

Increased competition for resources is a primary cost of group-living (Crook and Gartlan 1966, Alexander 1974, Wrangham *et al*. 1993, Chapman *et al*. 1995, Wheeler *et al*. 2013) and such competition may take the form of contests between individuals (e.g., Caraco and Wolf 1975, Janson 1985, Holekamp *et al*. 1996). However, resolved, differentiated dominance relationships, often associated with strong within-group contest competition (van Schaik 1989, Koenig *et al*. 2013, Pruetz and Isbell 2000), may help lower conflict-related costs (Bernstein 1981). These aspects of dominance relationships have attracted sustained research interest over the last hundred years (Hobson 2022). Dominance structure in animal societies ranges from being egalitarian, wherein the competitors have similar chances of winning, to being highly differentiated and linearly ordered, wherein predictability of contest outcomes facilitates resolved dominance order. In the latter structure, higher rank often confers priority of access to resources that may enhance the individual’s reproductive success (e.g., Janson 1985, van Schaik 1989, Holekamp *et al*. 1996, Shivani *et al*. 2022).

Socioecological theory predicts the placement of female-bonded societies along the egalitarian- linear dominance continuum based on ecological factors (Wrangham 1980, van Schaik 1989, Isbell 1991, Sterck *et al*. 1997, reviewed in Koenig *et al*. 2013). This theory predicts predominantly within-group scramble competition when food is dispersed and cannot be usurped by individuals, leading to rare food-related agonistic contests and non-differentiated or egalitarian dominance relationships. In contrast, high within-group contest competition is expected if food distribution is clumped and high-quality feeding sites are usurpable within groups, promoting frequent food-related contests and strongly expressed, well-resolved dominance rank relationships (e.g., Janson 1985, Pruetz and Isbell 2000). Past efforts to understand the role of ecological differences in shaping sociality and dominance have concentrated on primates (reviewed in Koenig and Borries 2009, Clutton-Brock and Janson 2012) and largely on stable groups (but see Holekamp *et al*. 1996, Archie *et al*. 2006, Smith *et al*. 2008 for high fission-fusion species). However, within-group competition may be lower in species showing fission-fusion dynamics, in which animals have the flexibility to change group size and composition over the short term, and such fluid associations could reduce the costs of within-group competition (e.g., Smith *et al*. 2008, Lehman *et al*. 2006).

Female elephant social groups are a good system to study whether the predictions of socioecological theory hold in high fission-fusion species. Elephants occupy a variety of habitats and substantial intra- and inter-specific variation in group living has been found despite the small number of elephant populations studied so far. Groupings of one or more females and their offspring are found in all three elephant species – *Loxodonta africana*, *L. cyclotis*, and *Elephas maximus* – (Douglas Hamilton 1972, McKay 1973, Sukumar 1989, Turkalo and Fay 1995), but long-term monitoring of individually-identified elephants has revealed differences across species/populations (see Moss 1988, Wittemyer *et al*. 2005, Archie *et al*. 2006, de Silva *et al*. 2011, de Silva and Wittemyer 2012, Fishlock and Lee 2013, Turkalo *et al*. 2013, Nandini *et al*. 2017, 2018, see Athira and Vidya 2021). For instance, these species/populations differ in group (analogous to “party” in the primate literature) size and fluidity in associations among community members due to varying extents of fission-fusion dynamics. African savannah elephants form core social groupings of females (family groups or second-tier units), led by a matriarch (the oldest female in the group, Moss 1988), that form well-connected and hierarchically nested, multilevel female societies (comprising bond groups/kinship groups and clans, or third- and fourth-tier units; Douglas Hamilton 1972, Moss and Poole 1983, Wittemyer *et al*. 2005). In contrast, African forest elephants have highly fluid associations and much smaller groupings (Fishlock and Lee 2013, Turkalo *et al*. 2013, Schuttler *et al*. 2014). Female Asian elephants are organised into communities or clans (identified through social network modularity analysis) with less hierarchical social organization and weaker associations, and smaller group (party) sizes than those in African savannah elephants (de Silva and Wittemyer 2012, Nandini *et al*. 2018). Differences in group sizes and social structure were attributed to ecological differences, such as food resource availability and distribution, as well as predation risk (de Silva and Wittemyer 2012, Nandini *et al*. 2018). However, differences in social structure were also observed between the two populations of Asian elephants examined, despite some similarities between the study areas and similar group sizes (de Silva *et al*. 2011, Nandini *et al*. 2018). Thus, given the variability in female social structure between the few populations of the three elephant species studied so far, whether there is a corresponding difference in dominance structure demands attention.

In the African savannah elephant, despite fission-fusion dynamics and despite food resources being widely dispersed, strong female dominance structure was found within family groups (Archie *et al*. 2006: in the Amboseli population, Kenya, and the Tarangire population, Tanzania), as well as between matriarchs of family (second-tier) groups (Wittemyer and Getz 2007: in the Samburu population, Kenya). Age played an important role in shaping dominance, with older females having a strong advantage in both cases, and dominance ranks were age-ordered rather than being based on relatedness, demonstrating weak nepotistic bias in within-group dominance (Archie *et al*. 2006). Conflict within family groups was not lower among closer relatives and associates (Archie *et al*. 2006). A prominent role of matriarchs was found in influencing dominance relationships between non-matriarchal females from different family groups (Wittemyer and Getz 2007), adding to previous findings on the importance of matriarchs in female African savannah elephant societies (Moss 1988, McComb *et al*. 2001, 2011).

In comparison, in a study on Asian elephant, females showed a small number of agonistic interactions, weaker effects of age, and unresolved dominance (de Silva *et al*. 2017, based on the Uda Walawe population, Sri Lanka). However, dominance relationships have not been specifically characterised within the most inclusive female social units or ‘societies’ (*sensu* Struhsaker 1969, “upper level” in Grueter *et al*. 2020, see also Moffett 2025 pp. 5-10) so far in the Asian elephant, with dominance interactions within as well as between social units (communities) being combined (in addition to examining interactions between communities) in de Silva *et al*. (2017). A comparable knowledge of within-group dominance in different elephant species is, therefore, still missing. Thus, to develop our understanding of elephant socioecology, we present here, our study of agonistic behaviour and dominance relationships within multiple social groups of female Asian elephants in the Kabini population, southern India, which is part of the largest contiguous population of Asian elephants in the world.

In our study population of Asian elephants in Kabini, communities identified from female association networks (using modularity) represent the most inclusive social units and are termed clans (Nandini *et al*. 2018). Females show fluid associations within clans, forming groups (“parties”) whose compositions can change over several hours due to fission-fusion dynamics. Although clans are of different sizes (due to demography), the average adult female group sizes observed are small and similar across clans (2-3 adult females), suggesting constraints on group size (Nandini *et al*. 2017, reviewed in Jabili *et al*. 2025). This, and the frequent agonism in large groups (Gautam and Vidya 2023) suggested that within-group competition might be important in shaping relationships within clans, as argued in socioecological models (van Schaik 1989, Isbell 1991). Associations between females belonging to different clans are extremely rare, despite groups (parties) from multiple clans simultaneously visiting the small resource-rich grassland around Kabini (length <15 km, maximum width <2 km), suggesting that the rules of clan membership and association are well defined (Nandini *et al*. 2017, Gautam and Vidya 2023).

We examined whether female Asian elephants had differentiated within-clan dominance relationships, and whether the outcomes and rates of within-clan agonistic interactions were influenced by age and/or social proximity (extent of association) among clan members. Since age is correlated with body size (Sukumar *et al*. 1988), which, in turn, could be correlated with dominance outcome, we expected older females to win against younger females. However, despite this expected effect of age on dominance rank, because of fluid grouping and small group sizes, we did not expect the matriarch (oldest female in the clan) to be the most dominant or most central (in terms of associations) female in the clan necessarily. We expected the rate of agonistic interactions to be lower between dyads with larger age differences than in age-matched dyads because larger differences in age/size might preclude the need for overt agonism, whereas females more closely matched in size might have less resolved dominance relationships and enter contests with each other more frequently. We also expected lower rates of conflict between females that showed greater social proximity measured by association index (see Methods), because spending more time together might resolve dominance relationships and lower conflict.

## METHODS

### Field sampling

We conducted field sampling in Nagarahole and Bandipur National Parks and Tiger Reserves, southern India (see Nandini *et al*. 2017 for details of the study area). The two National Parks are separated by the Kabini reservoir created by the Beechanahalli dam built on the Kabini river. As the dry season (mid-December – early June) progresses, the receding backwaters forms a grassland habitat, providing water and food to elephants and other wildlife. We collected most of the behavioural data on dominance from this open grassland habitat as it offered high visibility. We carried out field work from March 2009 to June 2013, sampling between about 6:30 AM to 5:45-6:45 PM, depending on field permits. Female elephant groups (parties) were identified as a set of adult female elephants and their dependent offspring that showed coordinated movement (especially to or from a water source) or affiliative behaviour and were within ∼50–100 m of one another (Nandini *et al*. 2017). All elephants sighted were aged, sexed, and individually identified based on natural physical characteristics and a database built up as part of the long-term Kabini Elephant Project (see Vidya *et al*. 2014). Individuals were placed into age-classes based on skull size, body size, the top fold of the ear, and other body characteristics, using the Forest Department’s semi-captive elephants of known ages in the area as a reference, and approximate dates of birth were assigned (see Vidya *et al*. 2014). We refer to females 10 years of age (since they could often reproduce from that age) or older as adults (as in Nandini *et al*. 2017).

### Behaviour data collection

Data on agonistic interactions were obtained through *ad libitum* and focal group sampling (Altmann 1974). Agonistic behaviours such as charges, chases, pushes, shoves, displacements, supplants, etc., and subordinate behaviours such as avoidance were recorded (Supplement Table S1). All the focal sampling and most of the *ad libitum* sampling were video-recorded using a Sony HDR-XR100E video camera. Although we noted down agonistic interactions in the field, we subsequently scored all the video footage in the lab to confirm the behaviours, their outcomes, and the identities of animals engaged in agonistic interactions. In this paper, we report on interactions only between adult females (henceforth called females). Agonistic interactions between two females could include a single interaction at a time or a series of interactions. When there were multiple interactions between the same pair of females, only the first interaction was considered independent and the subsequent interactions were considered non-independent if they occurred within 15 minutes of the first (see Gautam and Vidya 2023). We recorded the winner and loser of each interaction, as well as the eventual winner and loser of each independent interaction (see below).

During an agonistic interaction, if an individual was able to make the other individual move from its feeding site or turn away, the former was considered the winner, and the latter, the loser. There was no winner if the recipient ignored the initiator or retaliated ineffectively. If the recipient retaliated in such a manner as to make the initiator move away, the recipient was considered the winner. An individual initiating a subordinate behaviour in the absence of another initiating agonism was considered the loser (Supplement Table S1).

### Data on female associations

We used previously collected long-term data (March 2009 to July 2014) on associations amongst females (Nandini *et al*. 2017) to quantify social proximity and centrality (see Data Analysis below). Females that were part of the same group were considered to associate together in that sighting.

### Data analysis

We included only independent agonistic interactions between females in all our analyses. We used information on the eventual winners and losers if there was a sequence of non-independent interactions.

### Dominance structure within clans

To analyse dominance structure within clans, we looked at the following aspects of orderliness in dominance relationships in five common clans: unidirectionality, linearity, steepness of dominance hierarchy, triangle transitivity, and triad motifs in the dominance network. Unidirectionality of dominance was examined by ruling out reciprocity in dominance relationships, using Mantel *Z* test of reciprocity and Hemelrijk’s *Rr* test of relative reciprocity (Hemelrijk 1990). These test the correlation between the dominance matrix and its transpose. Linearity, which tests for whether competitors can be ordered in a linear dominance hierarchy, was assessed using de Vries’ index *h*’ (de Vries 1995; modified Landau’s *h*). Linear dominance hierarchies are obtained when triadic relationships are transitive rather than circular.

Using data on wins and losses, we quantified the dominance score of females within their clan using the normalised modified David’s score (MDS; DS based on *D*ij, see de Vries *et al*. 2006, Supplement text), and the steepness of the dominance hierarchy (*k*) as the absolute slope of the regression of the normalised MDS of clan members on their ranks. David’s score (DS; David 1987, 1988) is based on the sum of dyadic proportions of wins against and sum of dyadic proportions of losses to different individuals, taking into account the latter’s dyadic proportions of wins and losses. To infer statistical inference, we compared the observed values of linearity, unidirectionality, and hierarchy steepness with their respective randomised values using 5000 permutations, and calculated the proportion of times the randomized value was greater than or equal to the observed value. These analyses were performed in SOCPROG 2.4 (Whitehead 2009) run on MATLAB 7 (2004).

We complemented the measurement of linearity above by using triangle transitivity since sparsely-filled dominance matrices may not uncover linearity (unobserved relations are filled in different ways when calculating indices of linearity). Triangle transitivity (*t*tri) is an index based on the proportion (out of all triangles) of transitive triads, P(*t*), wherein P(*t*)=*N*transitive / (*N*transitive + *N*cyclic) (Shizuka and McDonald 2012). Since P(*t*) of a random network is expected to be 0.75, *t*tri is calculated as 4(P(*t*) – 0.75) such that zero corresponds to a random network and one to a network with all the triangles being transitive and thus showing resolved dominance. We also conducted triad census of the dominance network (see Wasserman and Faust 1994) for each of the five common clans. Although the number of complete triads (with all three pairwise relationships known) could be small, incomplete triads that were double- dominants (A dominates B and C) or double-subordinates could be additionally used to infer transitivity (Shizuka and McDonald 2015, see de Silva *et al*. 2017). When dominance order is present, transitive and double-dominant motifs are more abundant than expected, whereas pass- alongs and cyclic triads are rarer than expected by chance. Triad census and transitivity calculation and their statistical significance were carried out by simulating 1100 dyad census- conditioned random graphs (see Supplement text) based on the code provided in Shizuka and McDonald (2012, pp. 933 and Supplement therein), using the *statnet* package (Handcock *et al*. 2019) in R.

### Effects of age and social proximity on agonism and dominance relationships

To examine whether age influenced the initiation or outcome of agonistic interactions, we used data from all observations (focal and *ad-libitum*) and calculated the percentages of interactions won by initiators and recipients and by older and younger females. We compared the ages (on the day of the interaction) of initiators and recipients, and of winners and losers, using Wilcoxon’s matched-pairs tests (in Statistica 7; StatSoft 2004). As mentioned in the previous section, we quantified the dominance of females within their clans using the normalised MDS. For each of the five commonly observed clans, we also regressed these dominance scores of females against age (estimated at the mid-point of the study period, as each female was represented only once in this analysis) and tested for statistical significance using permutation test in R (R Core Team 2018; see Supplement text).

To examine whether age and social proximity affected the rate of agonistic interactions, we prepared a matrix of the dyadic rates of agonistic interactions, i.e., the number of agonistic interactions per hour for pairs of females, based only on focal observations, for each of the five commonly observed clans (see also Supplement text). We included only those dyads for which we had total focal observations of almost an hour or more, to avoid inflated/deflated estimates of dyadic agonism rate due to short observation time. We carried out Mantel tests (Mantel 1967) within each of the five common clans to examine if the dyadic agonism rate matrix was correlated with matrices of age difference and social proximity of female dyads. Social proximity had previously been quantified by Nandini *et al*. (2017) as the Association Index (AI) between pairs of females based on long-term sighting data (see Supplement text) and we used these AI values. Mantel tests were performed using the package *vegan* (Dixon 2003), with 5000 permutations, in R (R Core Team 2018).

### Effect of age on females’ centrality in the social network

We used the social network built using the long-term data on female associations (Nandini *et al*. 2017, see Supplement text) and calculated three measures of centrality for females: a) degree centrality (the number of associates a focal female has, or the number of nodes a focal node is connected to in the network), b) closeness centrality (a measure of how close a female, or node in the network, is to others, calculated as the inverse of the sum of path lengths from a focal node to all the other nodes), and c) betweenness centrality (a measure of how important a female is in the connectedness of the network, calculated as the proportion of all shortest paths between all other pairs of nodes that pass through the focal node) (see Wasserman and Faust 1994). We calculated these centrality measures using Gephi 0.7 (Bastian *et al*. 2009). We then regressed these centrality measures of females on their age, separately for the five common clans in *R*. Similarly, we tested for the relationship between centrality and dominance scores (normalized MDS) of females in each clan. For all these tests involving individual attributes of females, we performed 1000 permutations in R to test for statistical significance (see Supplement text).

## RESULTS

We recorded a total of 794 (475 independent, 319 non-independent) within-clan agonistic interactions, of which 252 independent and 163 non-independent interactions were recorded during 113.78 hours of focal group observations and 223 independent and 156 non-independent interactions were recorded during *ad-libitum* sampling (Table S2).

### a) Dominance structure within clans

Unidirectionality in dominance relationships based on the lack of absolute reciprocity was found in all five clans, and unidirectionality based on the lack of relative reciprocity was found in four clans (significant relative reciprocity in Lisa’s clan) (Figure 1, Table S3, age-ordered dominance matrices in Figure S1). Neither linearity based on de Vries’ *hꞌ* nor steepness of dominance hierarchy was statistically significant (Figure 1), but it should be noted that this likely resulted from the high sparseness of dominance matrices (dominance relationships were not known for ≥60% of all dyads in four out of five clans, see Table S3).

**Figure 1.**
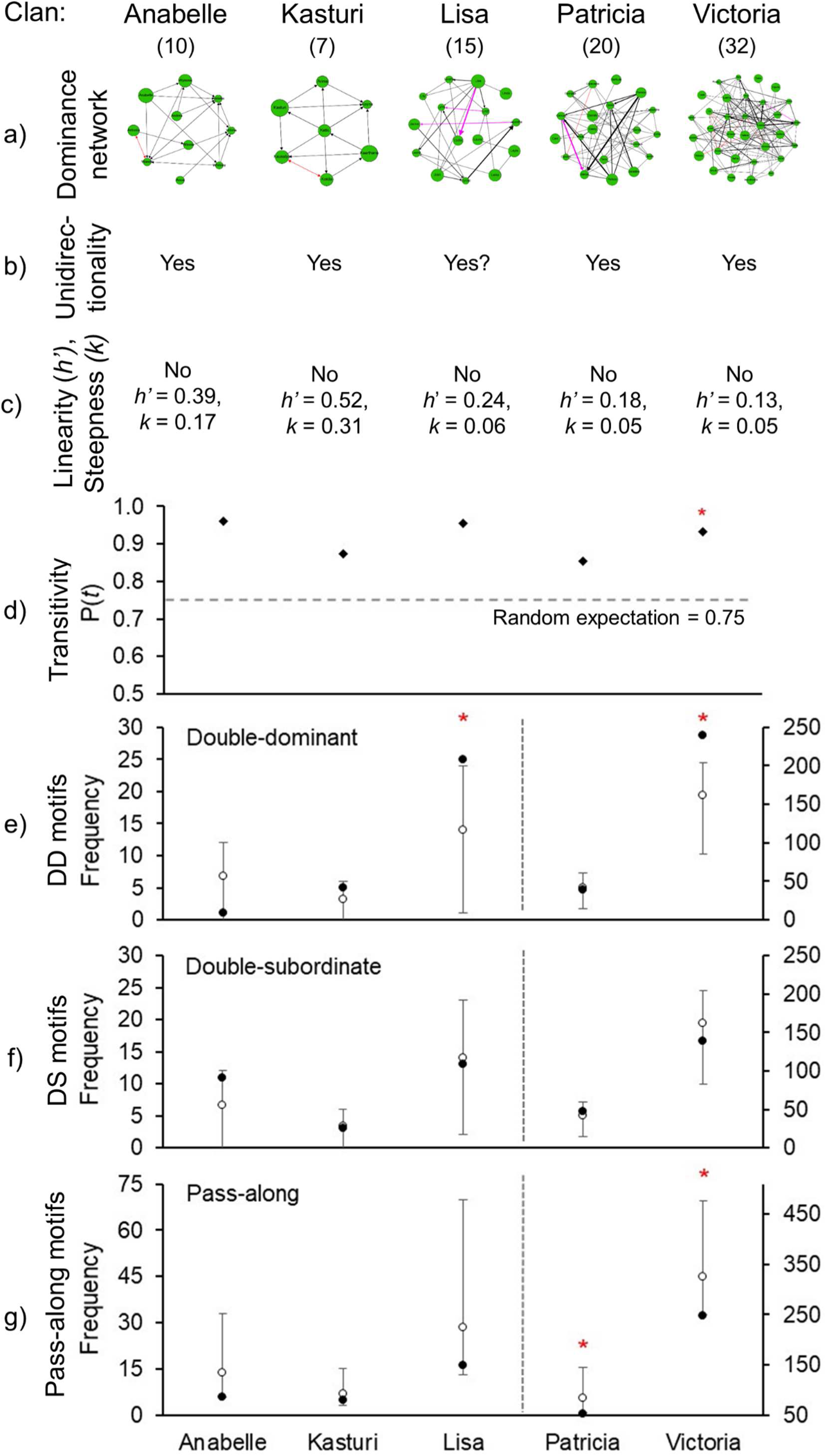
Dominance structure in five focal clans. a) dominance networks of females (shown as nodes), with the node size representing age relative to other clan members and arrows showing the direction of agonistic interactions (networks elaborated in Supplement Figure S2), presence of directionality, c) linearity (based on de Vries’ 1995 h’) of dominance and steepness (k; see Table S3), d) proportion of transitive triads, P(t), in dominance networks, along with the random expectation (dashed line), and e) – g) triad motifs, with the observed (filled circles) and random (open circles are the means and bars are 95% of the distribution) numbers of double-dominants (DD), double-subordinates (DS), and pass-alongs shown. The vertical lines in e) – g) separate the clans whose values are plotted on the primary and second Y axes (as the frequencies are very different). Red asterisks mark statistically significant values in d) – g).

Analysis of dominance order using triad census and triad transitivity of dominance networks suggested the presence of orderliness in some clans. The proportion of transitive triads (P(*t*)) in all five focal clans was greater than 0.75 (the random expected value), but the difference between the observed and random P(*t*) values was statistically significant in only Victoria’s clan (Figure 1d, Supplement Table S4, dominance networks elaborated in Figure S2). The observed numbers of double-dominant triads were significantly higher than random (observed values greater than 95% of the respective random values) in Lisa and Victoria’s clans, and the numbers of pass-along triads were significantly lower than random expectation in Patricia and Victoria’s clans (Figure 1, Table S4). The observed numbers of double-subordinate triads were not significantly different from random expectation in any clan.

### b) Effects of age and social proximity on agonism and dominance relationships

Almost all (98%) the independent agonistic interactions had clear outcomes, with the initiators winning over 95% of the resolved interactions, and a majority (75%) of the independent interactions being initiated by older individuals (summarised in Supplement text and Table S5). Overall, initiators were older than recipients (Wilcoxon matched pairs test: *Z*=12.176, *P*<0.001, *N*=475) and winners were older (average ± SD = 36.4 ± 14.56 years) than losers (average ± SD = 23.7 ± 12.42 years) (Wilcoxon’s matched-pairs test: *Z*=12.977, *P*<0.001, *N*=465 interactions) (Figure 2, see Supplement text, Table S5). Older females won in 76% of the independent interactions with clear outcomes, whereas reversals against older females were seen in the remaining 24% (Supplement text, Supplement Table S6). This pattern was not due to the age- structure of the interacting females (see Supplement text, Figure 2d).

**Figure 2.**
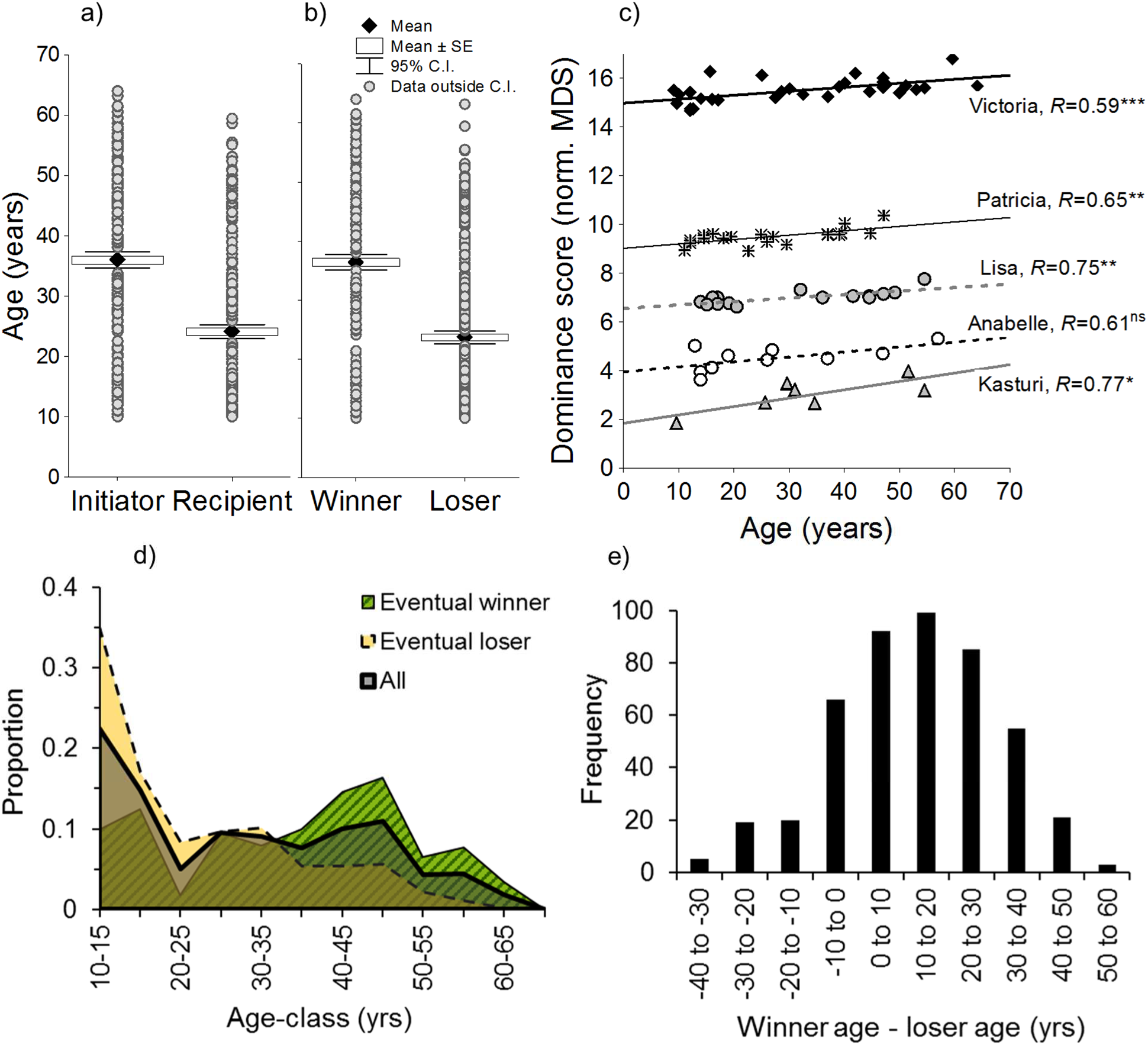
Effect of age on dominance. a) Initiators were older than recipients and b) winners were older than losers, in independent dominance interactions. c) Older females tended to have higher dominance scores (ns: R not significant, *: P<0.05, **: P<0.01, ***: P<0.001) (dominance scores are to be compared only within clans as MDS values are influenced by the number of females involved in dominance in each clan). d) The proportion of females of different age-classes from all clans who participated in agonistic interactions (all; thick black line), and proportions who were winners (solid line with hatched area) and losers (dashed line) (ages corresponding to all 465 resolved agonistic interactions are included here). e) Age difference between winners and losers based on all independent interactions.

Older females had a tendency to have higher dominance scores, with statistically significant regressions in four out of five clans (Figure 2c, Supplement Table S7, age-ordered dominance matrices in Figure S1). The percentage of age-reversals was 35% in Anabelle’s clan that did not show a significant regression of age on dominance score, and ranged from 23-26% in the other four common clans (Supplement Table S6). Despite the overall effect of age on dominance score, the matriarch was not the most dominant female in two (Victoria’s, Kasturi’s) of the four clans with significant regressions (Figure 2c). Contrary to expectation, the dyadic rate of agonism within clans was not significantly correlated with the age difference within the interacting dyad in any focal clan (Figure 3, Table S10). However, the absolute age difference between the winner and loser was higher when the winner was older than the loser compared to when the winner was younger than the loser (age-reversal) (see Table S6, also Figure 2e).

**Figure 3.**
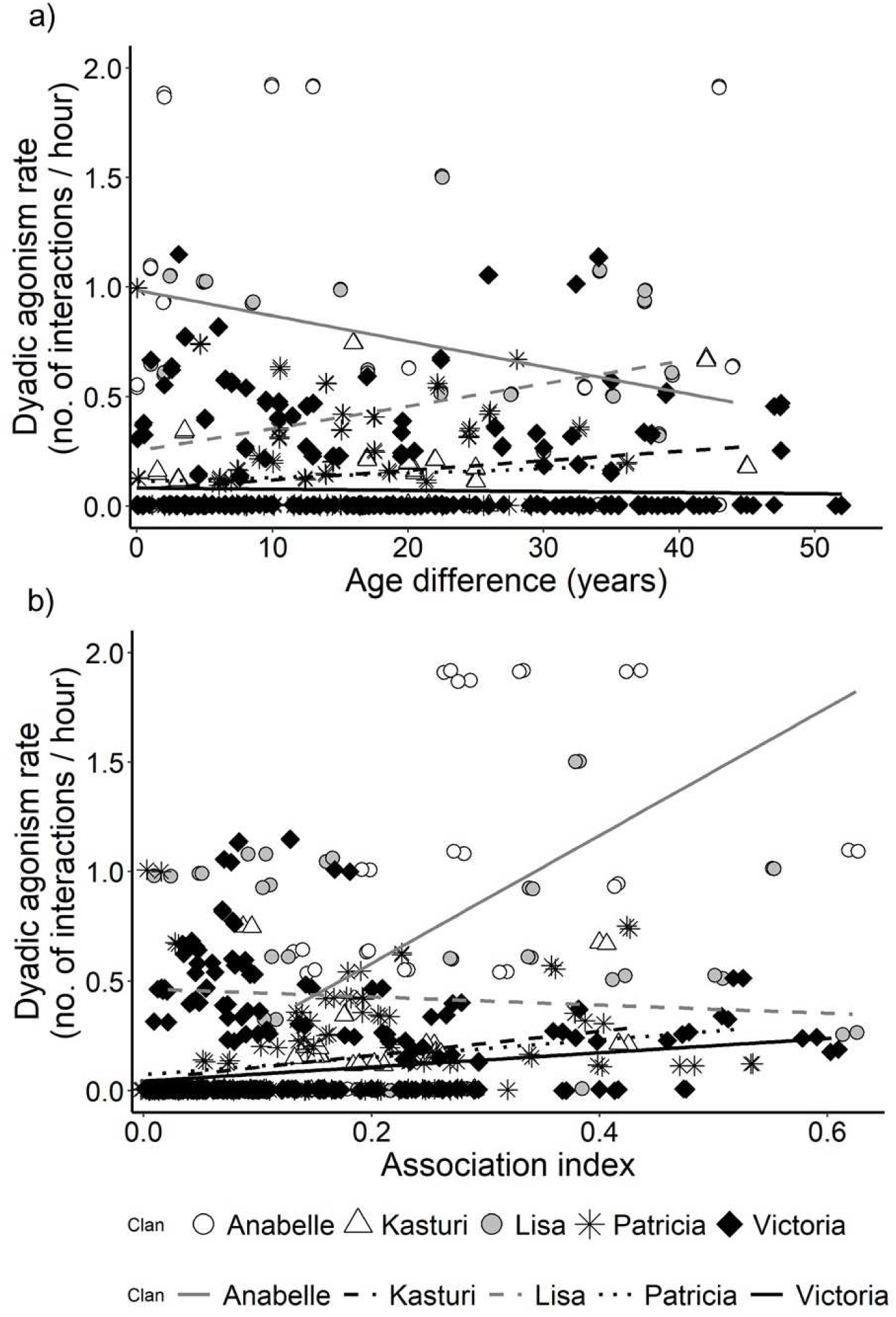
Association between the rate of dyadic agonism (i.e., between pairs of females shown as datapoints) and a) age difference and b) association index in five common clans. Based on Mantel tests, dyadic agonism was a) not correlated significantly with age difference in any clan, and b) positively correlated with association index in Anabelle’s, Patricia’s and Victoria’s clans. For better visualization, one dyad with an outlier agonism rate and two dyads with outlier association index values are not visible due to axis truncation in plots a and b, respectively (but included in Mantel tests).

The dyadic rate of agonism within clans was positively, albeit weakly, correlated with AI (social proximity) in three (Anabelle, Patricia and Victoria’s) clans (Figure 3, Table S10).

### c) Influence of age and dominance status on centrality of females in the social network

Female age did not predict degree centrality, closeness centrality, or betweenness centrality in any of the five common clans (Figure 4, Supplement Table S8). The matriarch (clan’s oldest female) was not more central than other clan members. Dominance score was also not significantly related to centrality attributes of females in any clan (Supplement Table S9).

**Figure 4.**
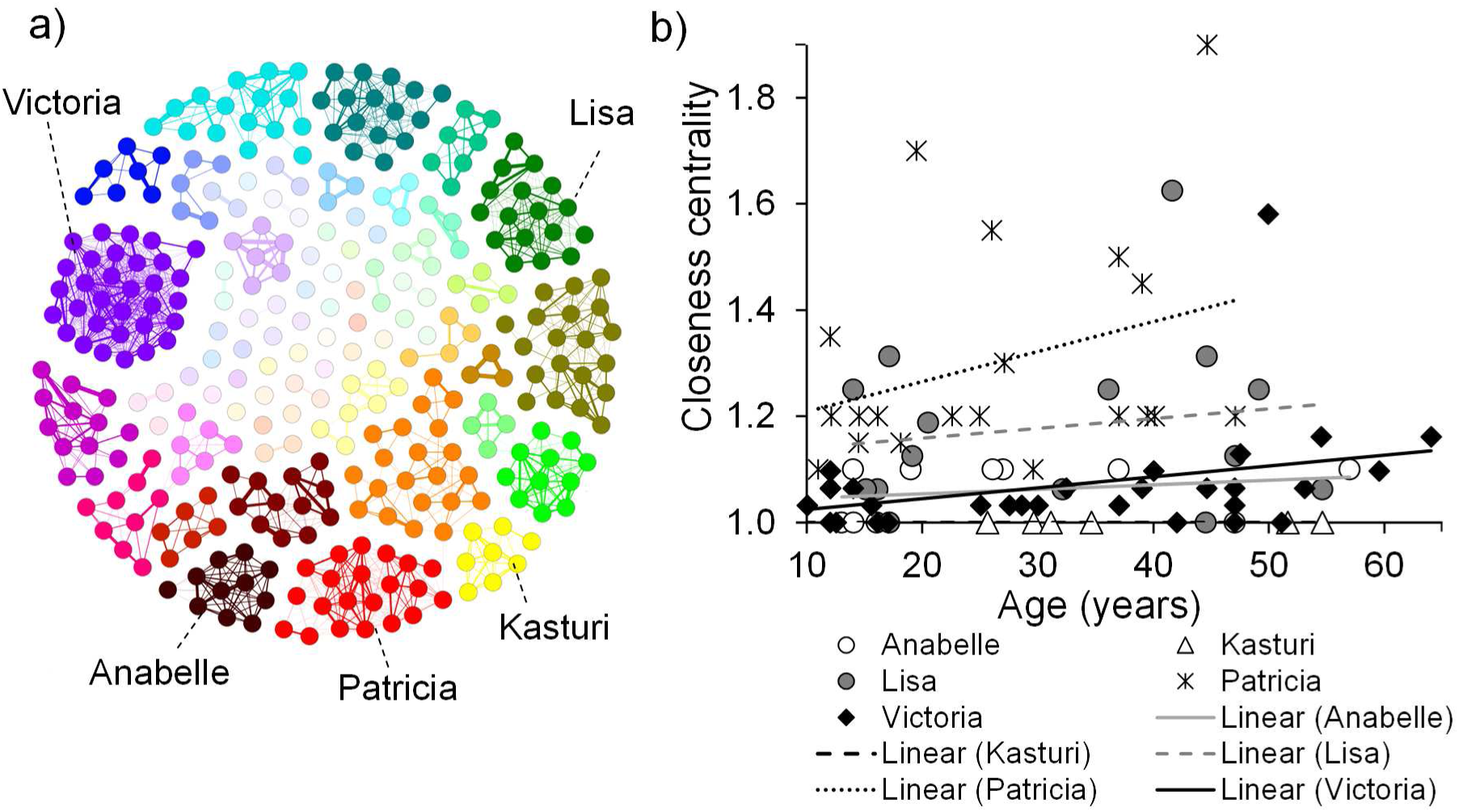
Relationship between age and centrality of females in the social network. a) Social network built using over five years of data on associations among females (represented by nodes), reproduced from Nandini et al. (2018; Group size differences may mask underlying similarities in social structure: a comparison of female elephant societies. Behavioural Ecology, 29, 145-159) by permission of Oxford University Press and focal clan names added. b) Scatterplots of closeness centrality (calculated from the association network) and female age in the five common clans.

## DISCUSSION

We conducted this study to understand agonistic and dominance relationships within clans, the most inclusive female social units in Asian elephants that show high fission-fusion characterised by changes in associations across hours (Nandini *et al*. 2018). We found dominance order based on age in these clans, but somewhat weak dominance structure in terms of transitivity in triadic motifs and percentage of reversals against age. Contrary to expectation, the age difference between clan members did not influence within-clan conflict as measured by the dyadic rate of agonism. Social proximity among clan members, measured as association index, also did not lower within-group conflict. The matriarch – the oldest female in the clan – was not necessarily the most dominant female, nor was she the most central in the social network based on long-term associations. This female dominance structure in Asian elephants is in contrast to the stronger dominance structure in African savannah elephants, as discussed below.

### Weaker dominance structure within female Asian elephant than African savannah elephant societies

All five focal clans in our study showed significant unidirectionality by at least one test, implying resolved dominance at the dyadic level. We found age to significantly affect contest outcomes. However, at the triadic level, we found mixed evidence of dominance resolution because although the observed proportion of transitive triads was greater than 0.75 in all the clans, only some clans showed statistically significant triangle transitivity (Figure 1d), greater abundance of triad motifs that lead to transitivity (Figure 1e-f) or statistically lower frequency of motifs leading to cyclicity (Fig. 1g) than expected by chance. Triad motif structures indicative of resolved dominance are common in animal groups (Shizuka and McDonald 2015 pp. 4) and our findings suggest somewhat weakly resolved dominance structure. We did not detect linearity or high steepness of dominance hierarchy using MDS, but it must be noted that our dominance matrices were too sparse (≥60% unknown relationships in four out of five common clans) to reliably detect statistical significance in these measures (see de Vries 1995, Saccà *et al*. 2022).

In contrast to our findings, stronger dominance structure had been found within family groups in African savannah elephants in Amboseli and Tarangire (Archie *et al*. 2006). Our study clans showed lower transitivity (average ttri=0.662, Table S4) than that seen in Amboseli (no cyclicity i.e., ttri=1.0, based on Archie *et al*. 2006, see Supplement text). Asian elephant clans also showed a weaker effect of age on dominance as reversals against older females were more frequent (23-35%) than in savannah elephants (4-6%, Archie *et al*. 2006). Significant linearity and transitivity had been seen among the matriarchs of second-tier family groups also in savannah elephants (Wittemyer and Getz 2007), but Asian elephants do not show similarly clear second-tier groups within clans.

Linearity and steepness were not significant within family groups in Amboseli either due to insufficient interactions observed (see Supplement text, Archie *et al*. 2006, Lee 1987). The average steepness (0.284) based on data from Amboseli (Archie *et al*. 2006, see Supplement text) was higher than that based on our focal clans (average=0.126, Table S3), but since steepness based on MDS decreases with an increasing proportion of missing relationships (Saccà *et al*. 2022), and this proportion was higher in our study clans than in family groups in Amboseli, how different the steepness values are actually is not clear.

According to socioecological theory, distinct dominance structures can arise under contrasting distributions of contestable, high-quality foods (van Schaik 1989, Sterck *et al*. 1999). Archie *et al*. (2006) suggested that African savannah elephants may be competing for rare high-quality foods (e.g., *Acacia* trees) and other resources (rubbing posts, water and mineral resources), which was also reported from Samburu, Kenya (Wittemyer and Getz 2007). In contrast, dominance was weak in our study despite mass flowering of bamboo, a clumped high-quality food, during the study period. While elephants fed on bamboo for considerable time, they showed agonism largely while feeding on grass, whereas contests for bamboo were rare (personal observations NS & TNCV, 2011-2013). Among other explanations, in their comparison of dominance between African savannah (Samburu) and Asian elephants (Uda Walawe, Sri Lanka), de Silva *et al*. (2017) suggested that greater fission-fusion tendency and greater rainfall in Asian habitats might lower competition and weaken dominance. We think that fission-fusion may not be a sufficient explanation for within-species differences in competition as we found more frequent agonistic interactions in our study population (Supplement text) than in Uda Walawe, despite possibly similar habitat conditions (see also Gautam and Vidya 2023). However, weaker dominance in Asian elephants compared to African savannah elephants could be plausibly explained by greater fission-fusion and, possibly, lower predation risk (that can reduce competition through decrease in group size and social cohesiveness) experienced by Asian elephants, which should be examined by future studies. It is interesting to note that Kabini was intermediate in social cohesiveness between Uda Walawe and Samburu (Nandini *et al*. 2018) and dominance structure/rate of agonism seems to mirror this pattern (while de Silva *et al*. 2017 combined agonistic interactions between and within social units (communities), the number of agonistic interactions within communities was small). Finally, variation in age-structure may also influence dominance. A visual comparison of age-structure of participating females in Kabini and Uda Walawe suggests that the youngest age-class (10-20 years, also the least dominant) is relatively more abundant in Kabini (Figure 2) than in Uda Walawe (Figure 2 in de Silva *et al*. 2017), which could have enhanced dominance resolution in Kabini. We did not have similar information for Amboseli for comparison.

While our inference of weak dominance in Asian elephants is consistent with de Silva *et al*.’s (2017) inference based on the Uda Walawe population, it should be noted that these inferences are based on different types of dominance networks. de Silva *et al*. (2017) did not specifically study dominance structure within social units and instead combined interactions within and between social units (analogous to between clans in Kabini) to build their dominance network. Since clans in Asian elephants are distinct from each other as associations between them are very rare (Nandini *et al*. 2018) and between-clan conflict is high (Gautam and Vidya 2023), analysing dominance within rather than between such communities is more informative of the social structure within ‘societies’ in the classical sense (*sensu* Struhsaker 1969). For an operational definition of the “most inclusive society” that is comparable across elephant species, we consider the “upper level” (just below the apex level that is connected only due to shared space and not sociality) in Grueter *et al*. (2020) to be particularly appropriate (see also Grueter and Swedell 2025 and Moffett 2025 pp. 5-10). This would correspond to *clan* in Asian and possibly *bond-group* or fourth-tier group (Wittemyer *et al*. 2005) in African savannah elephants.

### Age difference and social proximity do not reduce conflict or agonism

We found that initiators almost always won their interactions, suggesting that initiators can reliably assess their chances of winning. The basis for such assessment could be cues such as opponent age/size, combined with past experience of interactions. Indeed, older females initiated and won the majority of interactions. We also expected from age-based dominance asymmetry that the rate of agonistic interactions would be lower between dyads with larger age differences than in age-matched dyads as the latter might have less resolved dominance relationships and enter contests with each other more frequently whereas larger differences in age/size might preclude the need for overt agonism. The lack of this pattern seems to result from older females engaging in (and winning) contests with young females, as well as reversals against age-order. It is possible that these results are at least in part due to the rich resource habitat in which agonism was observed during the otherwise resource-poor dry season. Age-based dominance order would also suggest that younger females might avoid conflict with older clan members. This might indeed be the case: while “avoidance” as agonistic behaviour was rare (2-3% interactions were AVO or ADB, see Table S1 for description), most such instances (10 out of 13) involved younger females avoiding older females. Feeding away from older individuals may be another way for younger females to avoid conflict and this could be explored in the future.

We also expected closely associated females to show lower rates of agonism since such conflict is expected to decrease when individuals spend more time together. Contrarily, we found slightly higher levels of agonism among more closely associated females in three out of the five clans. This correlation, albeit with weak effect, is puzzling but may result if females who associate more frequently also forage in closer proximity, which then increases the probability of agonism compared to females in the same group who forage further apart. Examining the link between spatial proximity, obtained from individual positions during foraging, and agonism rates or association index could provide more insight into the effect of social proximity, as well as age difference, on agonism. Further, if females who are spatially close in the group are also close relatives, there might be advantages of such spatial structuring, which offset increased agonism; agonism within clans is quickly resolved and not intense based on the behaviours we find (Table S1). In African savannah elephants also, greater agonism was reported among more closely associated females who were also closer relatives, suggesting the absence of nepotistic bias in dominance within family groups (Archie *et al*. 2006; however, female dominance rank within the more inclusive social units (bond groups) was influenced by the matriarch’s dominance rank - Wittemyer and Getz 2007). Such role of kinship on dominance is yet to be examined in Asian elephants.

### Matriarch’s less prominent position in Asian elephant societies?

The matriarch’s role is prominent in African savannah elephant female groups, as matriarchs are repositories of socially and ecologically useful knowledge, and also influence the dominance status of their family members during conflict with other family groups (Dublin 1983, McComb *et al*. 2001, 2011, Wittemyer and Getz 2007, Mutinda *et al*. 2011). We found that while older females had higher dominance scores, in keeping with the significant proportion of reversals, the matriarch (oldest female in the clan) was not necessarily the most dominant female in her clan. The matriarch was the most dominant female in three out of five clans (and two of the four that showed significant correlations between dominance score and age). Moreover, the non-centrality of matriarchs in the association networks (Table S9) suggests that sociality may be not be centralized around matriarchs in the Asian elephant. It is not clear if these are different from corresponding values in the African savannah elephant as the oldest female was the most dominant female in seven out of 10 family groups in Amboseli (based on MDS, data from Archie *et al*. 2006). Centrality of matriarchs in social networks has not been reported. Matriarchs are likely to be repositories of knowledge in the Asian elephant also, even if their dominance status and social centrality are different from those in the African savannah elephant. Differences may arise due to more fluid grouping and smaller group sizes, as well as more temporally predictable resource distribution over the short term, in the Asian elephant (see Nandini *et al*. 2017, 2018), and cooperative predators and unpredictable resource distribution in the African savannah elephant. It would be interesting to examine the relative importance of the matriarchs in the two species, as well as the relationship between dominance, social connectedness, and social importance, in the future.

In conclusion, our study shows that female Asian elephant clans have age-based order, with advantages to older females, but dominance relationships are not strongly resolved. There is a significant proportion of reversals against age-order and the matriarch is not necessarily the most dominant or central female in the society. This contrasts with stronger, resolved dominance structure in the African savannah elephant and the keystone-like status of matriarchs. Several socioecological or demographic factors could have potentially shaped these differences. More in-depth studies on agonism and dominance, along with quantification of potential predictors like food distribution, predation risk, and individual attributes of females are required to understand mechanisms shaping dominance in elephant groups.

## ACKNOWLEDGEMENTS

This work was primarily funded by the Council of Scientific and Industrial Research, Government of India, under Grant No. 37(1375)/09/EMR-II and No. 37(1613)/13/EMR-II. National Geographic Society, USA, provided supplementary funds under Grant #8719-09 and #9378-13. We also thank the Department of Science and Technology, Government of India, for a Ramanujan Fellowship (to TNCV) under Grant No. SR/S2/RJN-25/2007 (dated 09/06/2008), through which TNCV was supported and some of the field work could be carried out. JNCASR provided logistic support. This work is part of the NRS’s doctoral thesis. We thank Carel van Schaik for comments on the thesis chapter version of this manuscript. NRS was supported as a Ph.D. student by JNCASR, and KP was supported as a Ph.D. student by the Council of Scientific and Industrial Research. No. 09/733(0152)/2011-EMR-I. HG was supported as a Research Associate by JNCASR. HG acknowledges the support from Science and Engineering Research Board, Department of Science and Technology, Government of India (National Postdoctoral Fellowship, No. PDF/2021/002320) during the manuscript’s preparation. The funders had no role in study design, data collection and analysis, decision to publish, or preparation of the manuscript. We thank the office of the PCCF, Karnataka Forest Department, for research permits, and various officials, from the PCCF and APCCF, to the Conservators of Forests and Range Forest Officers, to the staff of Nagarahole and Bandipur National Parks for permits and support at the field site. We thank Deepika Prasad for help with initial field data collection. We thank Mr. Gunda, Mr. Rajesh, Mr. Ranga, Mr. Krishna, Mr. Binu, Mr. Althaf and others for providing field assistance. We thank Ajay Desai for useful discussions on elephants. We thank Anvitha S for help with data analysis in R.

## Author contributions

NS and TNCV designed the study, NS and PK, with help from others, collected field data, NS and HG analysed data with inputs from TNCV, HG and NS wrote the first draft of the manuscript, all authors read and revised the manuscript and approved it for submission.

## Supplementary Material

### Supplement text

As mentioned in Methods, we obtained data on agonistic behaviour through *ad libitum* and focal group sampling (Altmann 1974). We included various dominant and subordinate behaviours (listed in Table S1), but only among adult females (10 years or older on the day of the interaction; however, age on the mid-point of the dataset was used for analysis of age versus dominance score, as each female was represented only once in the analysis). When agonistic interactions between two females included a series of interactions, only the first interaction between the two females was considered independent, and the subsequent interactions were considered non-independent if they occurred within 15 minutes of the first (see Gautam and Vidya 2023). The number of independent and non-independent agonistic interactions observed in focal and *ad libitum* observations for each clan are given in Table S2. We included only independent agonistic interactions in our further analyses.

### Rates of agonism

The average dyadic rates of agonism in the five common clans are shown in Figure 3 in the main text and reported in Table S10. The overall average dyadic rate of agonism was 0.354 interactions/dyad/hour (SD=0.374, *N*=455 dyads from 5 common clans, see Table S10 for clan- wise averages). It should be noted that this dyadic rate of within-clan agonism in Table 1 is a different measure from the rate of individual-level within-clan agonism reported in another study from the Kabini Elephant Project (Gautam and Vidya 2023), which examined agonistic competition experienced by an average female for the years 2015 and 2016. That rate of individual-level within-clan agonism in Gautam and Vidya (2023) was calculated as the total agonism experienced (initiated and received) by an average female in a focal group observation per hour), as opposed to the number of interactions per dyad per hour presented in the main text of this paper. The average (± SD) rates of individual-level within-clan agonism (calculated in the same manner as in Gautam and Vidya 2023) for different clans in the current study were 3.253 (± 1.827, *N*=5 focals, Anabelle’s clan), 0.674 (± 1.062, *N*=18, Kasturi’s clan), 1.025 (± 1.324, *N*=33, Lisa’s clan), 0.992 (± 1.417, *N*=27, Patricia’s clan), and 0.831 (± 1.113, *N*=55, Victoria’s clan) interactions experienced per female per hour. The overall average (± SD) rate of individual-level within-clan agonism in the five focal clans in the present study was 1.022 (± 1.346) interactions experienced per female per hour (*N*=120 focal observations at least 15 minutes long, total duration=80.03 focal group hours), which is similar to the rate (1.152 interactions per female per hour) reported in Gautam and Vidya (2023).

### Within-clan dominance structure

As explained in the main text, we used five aspects of orderliness in dominance relationships – unidirectionality, linearity, steepness of dominance hierarchy, triangle transitivity, and triad motifs – to characterise within-clan dominance structure. Unidirectionality of dominance was examined by ruling out reciprocity in dominance relationships, by testing the correlation between the dominance matrix and its transpose. This was done using Mantel *Z* test of reciprocity and Hemelrijk’s *Rr* test of relative reciprocity (Hemelrijk 1990). Linearity, which tests for a linear dominance hierarchy, was assessed using de Vries’ index *h*’ (de Vries 1995; modified Landau’s *h*). Linear dominance hierarchies are uncovered when triadic relationships are transitive (if A dominates B and B dominates C, A also dominates C) rather than circular (A dominates B, B dominates C, and C dominates A). We calculated the steepness of the dominance hierarchy as the absolute slope of the regression of the normalised modified David’s score (MDS; DS based on *D*ij, see de Vries *et al*. 2006) of clan members on their ranks. David’s score (DS; David 1987, 1988) is based on the sum of dyadic proportions of wins against, and sum of dyadic proportions of losses to, different individuals, taking into account the latter’s dyadic proportions of wins and losses. MDS accounts for chance occurrence of the observed outcomes based on different numbers of interactions between dyads (de Vries *et al*. 2006). We normalised MDS as (MDS + *N*(*N*-1)/2)/*N*, where *N* is the number of individuals (see de Vries *et al*. 2006). To infer statistical inference, we compared the observed values of linearity, unidirectionality, and hierarchy steepness with their respective randomised values using 5000 permutations and calculated the proportion of times the randomized value was greater than or equal to the observed value. The above analyses were performed in SOCPROG 2.4 (Whitehead 2009).

While most of the analyses in the paper use the MDS to measure dominance score of females, it has been shown that the steepness calculated based on MDS becomes lower with an increase in the proportion of unknown relationships (Saccà *et al*. 2022). Instead, the Average Dominance Index (ADI), which is the average proportion of wins by an individual against its opponents, was found to have lower bias in the face of unknown relationships and was recommended for calculating steepness. We, therefore, also report the steepness of dominance hierarchy as the absolute slope of the regression of the normalised (by multiplying into *N*-1) ADI of clan members on their ranks (see Saccà *et al*. 2022) in Table S3. It should be noted, however, that the ADI increases with an increase in the proportion of unknown relationships and it is possible to get slopes slightly higher than 1 using ADI (but not when using MDS) when there are missing relationships (Saccà *et al*. 2022). Similarly, in addition to the analysis using MDS, the slope of the relationship between ADI and age-based rank is shown in Table S7.

We complemented the measurement of linearity above by using triangle transitivity since sparsely-filled dominance matrices may not uncover linearity (unobserved relations are filled in different ways when calculating indices of linearity). Triangle transitivity (*t*tri) is an index based on the proportion (out of all triangles) of transitive triads, P(*t*), wherein P(*t*)=*N*transitive / (*N*transitive + *N*cyclic) (Shizuka and McDonald 2012). Since P(*t*) of a random network is expected to be 0.75, *t*tri is calculated as 4(P(*t*) – 0.75) such that zero corresponds to a random network and one to a network with all the triangles being transitive. We calculated *t*tri using the code of Shizuka and McDonald (2012, pp. 933) wherein the triads containing mutual ties (i.e., dyads with both members winning equally) were assigned weights based on the propensity of the triad to become transitive if one of its mutual dyads were to become clearly asymmetric. We wanted to test for statistical significance of the observed *t*tri by comparing it with *t*random that was obtained after simulating 1000 dyad census-conditioned random graphs (i.e., graphs with same number of participants and relationships but with a uniform probability for each individual winning over others). However, since some simulations would lead to undefined values of P*t* (in cases where *N*transitive + *N*cyclic was 0), resulting in fewer than 1000 valid values of randomized P*t*, we increased the number of simulations to 1100 for all clans so that inferences were always based on 1000 or more randomized values. The one-tailed *P* value was obtained as the proportion of times *t*random exceeded the observed *t*tri.

We also conducted triad census of the dominance network (Wasserman and Faust 1994), which examines different kinds of triad motifs, for each of the five common clans. When the number of triads with all three interactions is small, incomplete triads that include double-dominants (A dominates B, and A dominates C; B and C have not been seen to interact) or double- subordinates (B and C both dominate A; B and C have not been seen to interact) can be used to infer transitivity (Shizuka and McDonald 2015, see de Silva *et al*. 2017). In the case of double-dominants and double-subordinates, the triads will be transitive irrespective of the relationship between the dyad that is not observed to interact. On the other hand, when there are incomplete triads that show a pass-along relationship (C dominates A, and A dominates B; B and C have not been seen to interact), there is an equal probability of the triad becoming transitive or circular depending on the third relationship (if there is equal probability of the directionality of the third relationship). The observed numbers of these three triad motifs were compared with the respective values from 1100 simulations of random networks to test whether there were more double-dominants or double-subordinates than expected at random, and whether there were fewer pass-alongs than expected at random. The random networks had the same number of mutual, asymmetric, and null dyads as the observed network. Triad census and transitivity analyses were done based on the code provided in Shizuka and McDonald (2012), using the *statnet* package (R Core Team 2008) in R.

Results on within-clan dominance structure of the five common clans are presented in Tables S3 and S4 here and Figure 1 in the main text.

In the Discussion, hierarchy steepness, i.e., the slope of the relationship between MDS (normalized) and rank, observed based on our five common clans is compared with those of the ten family groups of African savannah elephants from the Amboseli population (Kenya), whose dominance data were reported in Archie *et al*. (2006). We used SOCPROG 2.4 to obtain hierarchy steepness based on the dominance matrices provided in Archie *et al*. (2006). Neither linearity (*h*’) nor steepness (based on *D*ij) were statistically significant in these 10 groups, perhaps due to matrix sparseness. The observed values of hierarchy steepness (based on MDS) in different family groups of African savannah elephants were: 0.183 (family group: AA), 0.562 (FB), 0.539 (CB), 0.148 (GB), 0.541 (DB), 0.220 (JA), 0.187 (EA), 0.133 (OA), 0.233 (EB), 0.090 (PC). Hierarchy steepness observed in Amboseli (average=0.284, SD=0.187) was over twice the steepness in Asian elephant clans from Kabini (average=0.126, SD=0.112). However, since steepness based on MDS decreases with an increasing proportion of missing relationships (Saccà *et al*. 2022), how different the steepness values are is not clear. Nevertheless, African savannah elephants in Amboseli seem to have a stronger dominance structure than Asian elephants in Kabini based on reversals and transitivity (only 4-6% reversals against age and 100% transitivity since there were no cyclic triads in Amboseli, Archie *et al*. 2006), as mentioned in the Discussion.

### Effect of age and social proximity on agonism and dominance relationships

We examined how age (on the day of the interaction) influenced initiation and outcome of independent within-clan agonistic interactions using data from all clans but conducted further detailed analyses using data from five common clans (Figure 2 in the main text). A summary of agonistic interactions and their outcomes is presented in Table S5 in this supplement. A majority (75%) of within-clan independent interactions involved initiation of agonism by individuals that were older than the recipients, and clear outcomes were seen in 98% of all interactions (see Table S5). Initiators were significantly older (average ± SD = 36.0 ± 14.59 years) than recipients (average ± SD = 24.2 ± 12.83) (Wilcoxon’s matched-pairs test: *T*=20078.50, *Z*=12.176, *P*<0.001, *N*=475 independent interactions) and winners were significantly older (average ± SD = 36.4 ± 14.56 years) than losers (average ± SD = 23.7 ± 12.42 years) (Wilcoxon’s matched-pairs test: *T*=16549.00, *Z*=12.977, *P*<0.001, *N*=465 independent interactions with clear outcomes). These ages did not change appreciably when unique dyads that interacted were used only once rather than each independent interaction being used (average ± SD: initiators: 35.4 ± 14.62 years, recipients: 24.2 ± 13.34 years, *N*=297 interacting unique dyads; winners: 35.7 ± 14.65 years, losers: 23.8 ± 13.01 years, *N*=291 interacting unique dyads with clear outcomes). In order to rule out the effect of the age-structure of the interacting animals on older females initiating / winning more interactions than younger females, we also compared the frequencies of winners in different age-classes with the expected frequencies of winners in different age-classes (if the interacting females – winners and losers – had equal chances of winning) using the *X*^2^ test. There was a significant difference between the observed and expected frequencies (*X*^2^=76.525, *df*=5, *P*<0.001, see Figure 2d in the main text; *X*^2^=42.226, *df*=5, *P*<0.001 if only unique dyads that interacted were used).

Reversals against age were seen in 24% of the interactions with clear outcomes (Supplement Table S6). The average difference in age between dyads in which younger individuals won against older individuals was lower than that of dyads in which older individuals won against younger individuals (Figure 2e, Supplement Table S6). We tested this by randomly assigning winner and loser identity for each interaction and calculating the average difference in age between winners and losers in each of 100 random datasets. The average difference in age between winner and loser when the winner was older (19.8 years, SD=12.65, *N*=355) was significantly higher than that based on 100 randomisations (*P* calculated as the number of random values>observed value / number of randomisations was <0.001). Similarly, the average difference in age between winner and loser when the winner was younger (10.1 years, SD=9.76, *N*=110) was significantly lower than that based on 100 randomisations (*P*<0.001).

This was seen in all five focal clans, although statistical significance (based on Mann-Whitney *U* test) was not observed in two clans when only the unique dyads were compared due to small sample size. While the smaller age differences could be attributed to errors in aging, about one-third to over half the unique dyadic age differences were greater than ten years (which would not result from such error) when younger individuals won, and about two-thirds to over 80% of the unique dyadic age differences were greater than ten years when older individuals won.

We used the data on wins and losses from five commonly observed clans, and quantified the dominance score of females within their clan as the modified David’s Score normalized by group size and number of interactions (de Vries *et al*. 2006), henceforth called normalized MDS, using SOCPROG 2.4 (Whitehead 2009) run on MATLAB 7 (MATLAB 2004). We also regressed the dominance score on female age (estimated at the mid-point of the study period – 29-April-2011 – as each female was represented only once in this analysis), separately in each clan, and found positive correlation coefficients. Statistical significance was tested by permuting age 1000 times and repeating regressions on the permuted data using R (R Core Team 2018, *lm* for regression). The results are presented in Figure 2c in the main text and Table S7.

If older females were more dominant, we expected lower rates of conflict in dyads in which the age difference was larger, because younger females might avoid conflict with older, dominant females. We, therefore, calculated the dyadic rate of agonism for each female dyad within each of the five commonly observed clans, as the number of agonistic interactions observed between females of the dyad per hour of focal observation of the dyad. Only those dyads that were observed for almost an hour or more were included to calculate the dyadic rate of agonism and only data from focal sampling were used for this. We carried out a Mantel test (Mantel 1967; Pearson’s correlation, 5000 permutations used in the package *vegan* (Dixon 2003) in R (R Core Team 2018) to examine the correlation between the dyadic agonism rate matrix and the corresponding age difference matrix in each of the five clans.

We similarly performed Mantel tests (Pearson’s correlation, 5000 permutations) to see if Association Index (AI) was related to dyadic agonism rate as social proximity might help in lowering conflict between females. We used previously calculated AI between females based on long-term data (March 2009 to July 2014) on associations (Nandini *et al*. 2017). In Nandini *et al*.’s (2017) study, groups had been defined in the same manner as the present study (coordinated movement and/or affiliative behaviour and females usually within 50-100 m of each another), and complete sightings with all the females identified had been used to build a social network. AI between pairs of females had been calculated as the Simple Ratio Index, such that AIAB = *N*AB/(*N*-*D*) where *N*AB was the number times females A and B were seen together (i.e., in the same sighting or group), *N* was the total number of sightings, and *D* the number of times neither A nor B was seen (Ginsberg and Young 1992). The results of the Mantel tests are presented in Table S10 here and in Figure 3 in the main text.

### Effect of age on females’ centrality in the social network

Using the above-mentioned social network built from female association data, we quantified centrality of each female to examine whether older females were more central in the association network than younger females. We quantified three measures of centrality: a) degree centrality (the number of associates of a focal female, or the number of nodes a focal node is connected to in the network), b) closeness centrality (a measure of how close a female, or node in the network, is to others, calculated as the inverse of the sum of path lengths from a focal node to all the other nodes), and c) betweenness centrality (a measure of how important a female is in the connectedness of the network, calculated as the proportion of all shortest paths between all other pairs of nodes that go through the focal node) (see Wasserman and Faust 1994, Borgatti 2006). We then performed regressions of centrality measures of females on age, separately for the five common clans, in R. To check if more dominant females were more central, we tested for the correlation between dominance scores (normalized MDS) of females and their centrality measures. Since network statistics of a node involve other nodes, making variates non-independent, we tested the significance of these regressions by permuting variable *X* 1000 times and repeating regressions on the permuted data using R (R Core Team 2018). The results are presented in Figure 4 of the main text and Tables S8 and S9 in this supplement.

*Note:* For analyses of centrality and dominance score in Victoria’s clan, the sample sizes differ slightly from those in Mantel tests of correlation between dyadic agonism rate, age difference, and association index. This was because i) one female’s (Farah) dominance score was undefined as she did not engage in any dominance, ii) one female (Faiza) was sighted in only *ad libitum* but not in focal observations (therefore, the dyadic rates of agonism could not be calculated for her), and iii) association data analysis in Nandini *et al*. (2018) had not included one young female (Floppy_ears). Because of these differences, the datasets on the four variables analysed differ with respect to the presence of Farah, Faiza and Floppy_ears as follows:

a. the dominance score matrix has 32 females and includes Faiza and Floppy_ears (for whom dominance interactions were observed), but it excludes Farah as her dominance score was not available since she did not interact in dominance.
b. the dyadic agonism rate matrix has 31 females and includes Floppy_ears and Farah, but excludes Faiza, for whom agonism rates were not available as she was not observed in focal observations used to calculate agonism rates.
c. the association index matrix has 31 females and includes Farah, but excludes Floppy_ears, who was not included in the analyses of association data in Nandini *et al*. (2018), and Faiza, as she was not present in the agonism rate matrix (b), with which the association index matrix is being correlated in this paper.
d. the centrality matrix has 31 females and includes Faiza but excludes Floppy_ears, whose centrality attributes were not available as she was not included in the analyses of association data, and Farah, as she was not present in the dominance score matrix (a), with which the centrality matrix is being correlated.

## Part B: Supplementary Tables

**Table S1.**
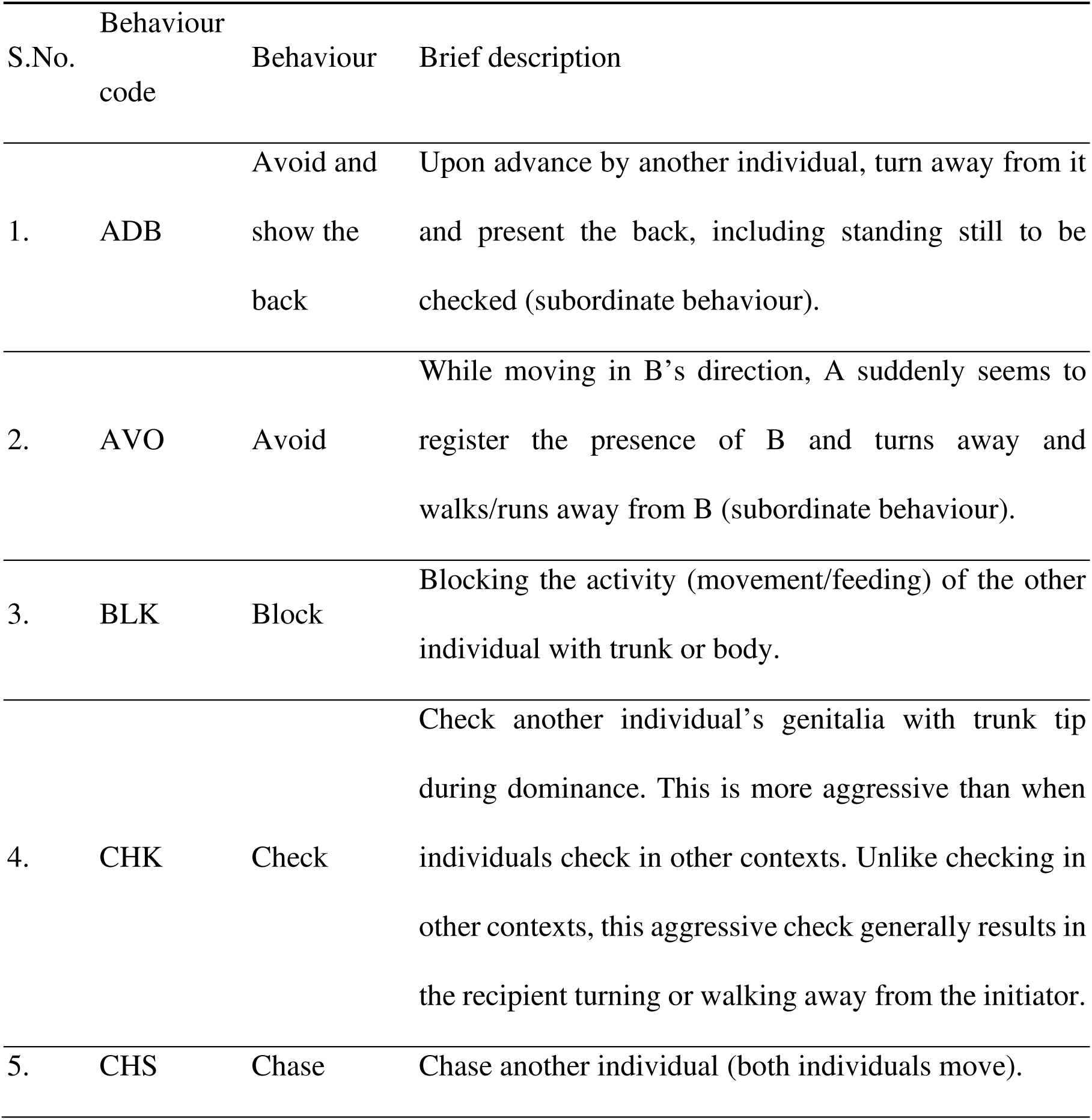
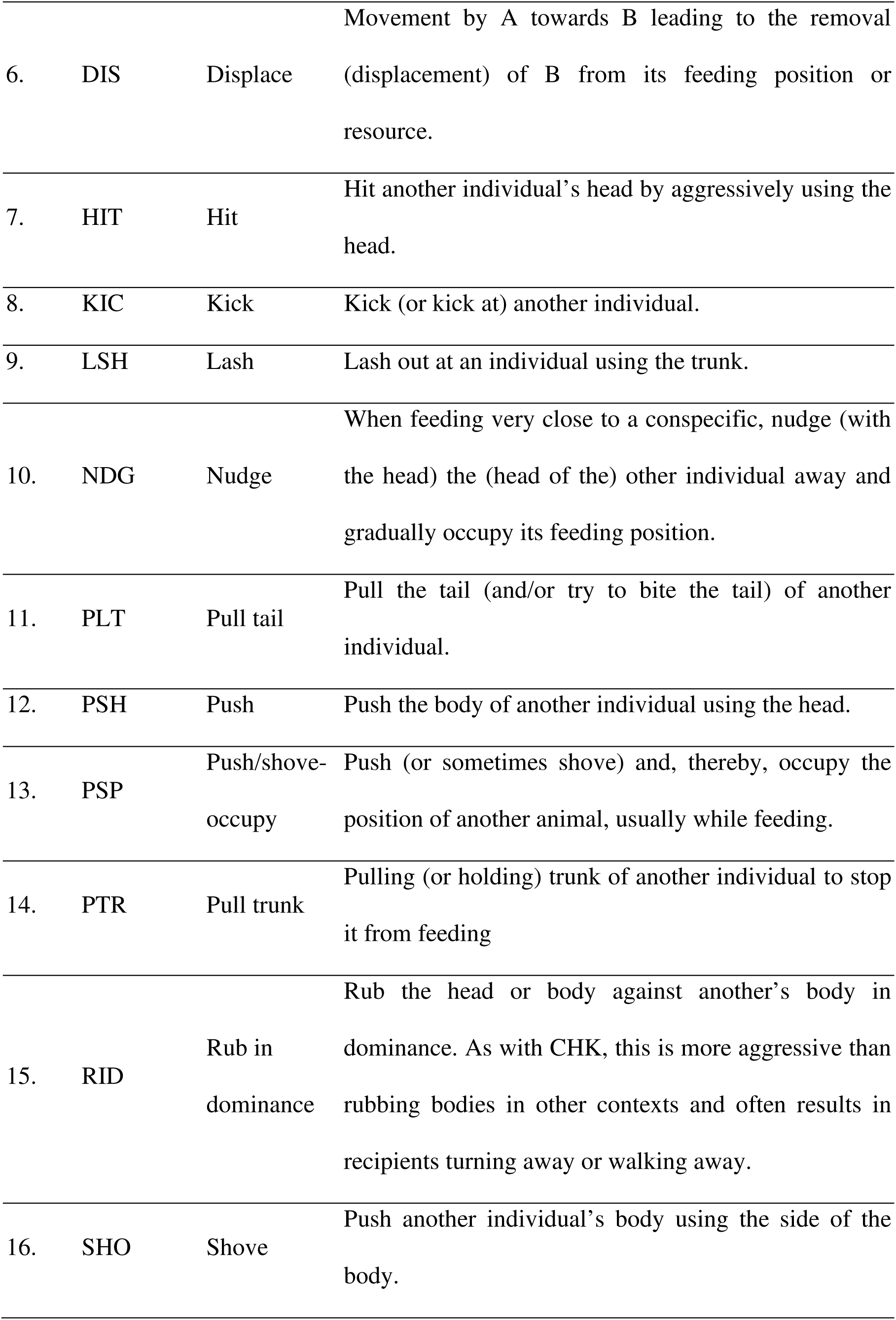
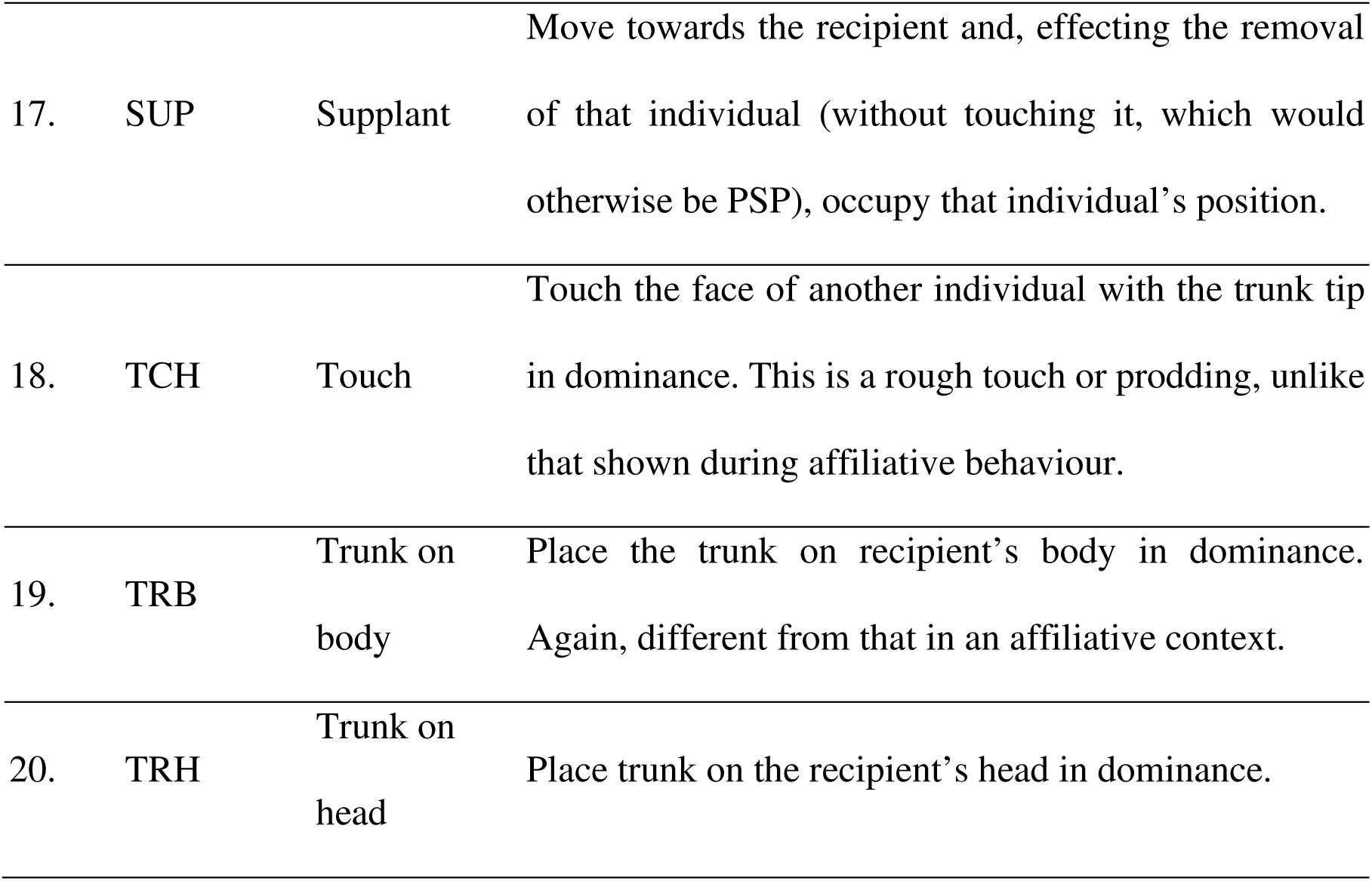
Different behaviours seen during agonistic interactions between females within clans. A and B are used as examples of initiator and recipient individuals. Subordinate behaviours are indicated within parentheses. This table is similar to supplements in Gautam and Vidya (2023), as many behaviour codes overlap.

**Table S2.** Summary statistics of within-clan agonistic interactions. Observations are divided into independent (I) and non-independent (NI) agonistic interactions and into focal and *ad-libitum* observations. Detailed analyses of agonistic relationships and dominance structure were done for the five common clans, whose rows are highlighted in grey.

| Clan name | Focal sampling |  |  |  | Ad-libitum sampling |  |  | Total |  |  |
| --- | --- | --- | --- | --- | --- | --- | --- | --- | --- | --- |
|  | Duration (hours) | No. of interactions |  |  | No. of interactions |  |  | No. of interactions |  |  |
|  |  | I | NI | Total | I | NI | Total | I | NI | Total |
| Anabelle | 2.58 | 32 | 12 | 44 | 5 | 0 | 5 | 37 | 12 | 49 |
| Andromeda | - | - | - | - | 1 | 1 | 2 | 1 | 1 | 2 |
| Fiola | 0.92 | 2 | 1 | 3 | 3 | 0 | 3 | 5 | 1 | 6 |
| Kasturi | 12.1 | 18 | 14 | 32 | 5 | 1 | 6 | 23 | 15 | 38 |
| Katrina | 0.47 | 1 | 0 | 1 | 4 | 0 | 4 | 5 | 0 | 5 |
| Lisa | 9.82 | 37 | 20 | 57 | 12 | 5 | 17 | 49 | 25 | 74 |
| Manasi | 1.5 | 3 | 3 | 6 | 0 | 0 | 0 | 3 | 3 | 6 |
| Mayura | - | - | - | - | 1 | 0 | 1 | 1 | 0 | 1 |
| Menaka | 4.2 | 7 | 7 | 14 | 1 | 0 | 1 | 8 | 7 | 15 |
| Mridula | 0.1 | 0* | 2 | 2 | 2 | 0 | 2 | 2 | 2 | 4 |
| Nakshatra | 6.78 | 7 | 6 | 13 | 8 | 7 | 15 | 15 | 13 | 28 |
| Olympia | 1.18 | 0 | 0 | 0 | 1 | 0 | 1 | 1 | 0 | 1 |
| Osanna | 1.75 | 1 | 0 | 1 | 2 | 0 | 2 | 3 | 0 | 3 |
| Patricia | 24.1 | 53 | 44 | 97 | 38 | 48 | 86 | 91 | 92 | 183 |
| Peggy | 2.18 | 0 | 0 | 0 | 2 | 0 | 2 | 2 | 0 | 2 |
| Thamarai | - | - | - | - | 1 | 0 | 1 | 1 | 0 | 1 |
| Tilottama | 0.9 | 3 | 2 | 5 | 2 | 1 | 3 | 5 | 3 | 8 |
| Unnati | 1.33 | 2 | 0 | 2 | 16 | 14 | 30 | 18 | 14 | 32 |
| Victoria | 43.9 | 86 | 52 | 138 | 118 | 75 | 193 | 204 | 127 | 331 |
| Yolanda | - | - | - | - | 1 | 4 | 5 | 1 | 4 | 5 |
| Total | 114 | 252 | 163 | 415 | 223 | 156 | 379 | 475 | 319 | 794 |

\*There are zero independent interactions here because the interaction began during *ad libitum* observations.

**Table S3.**
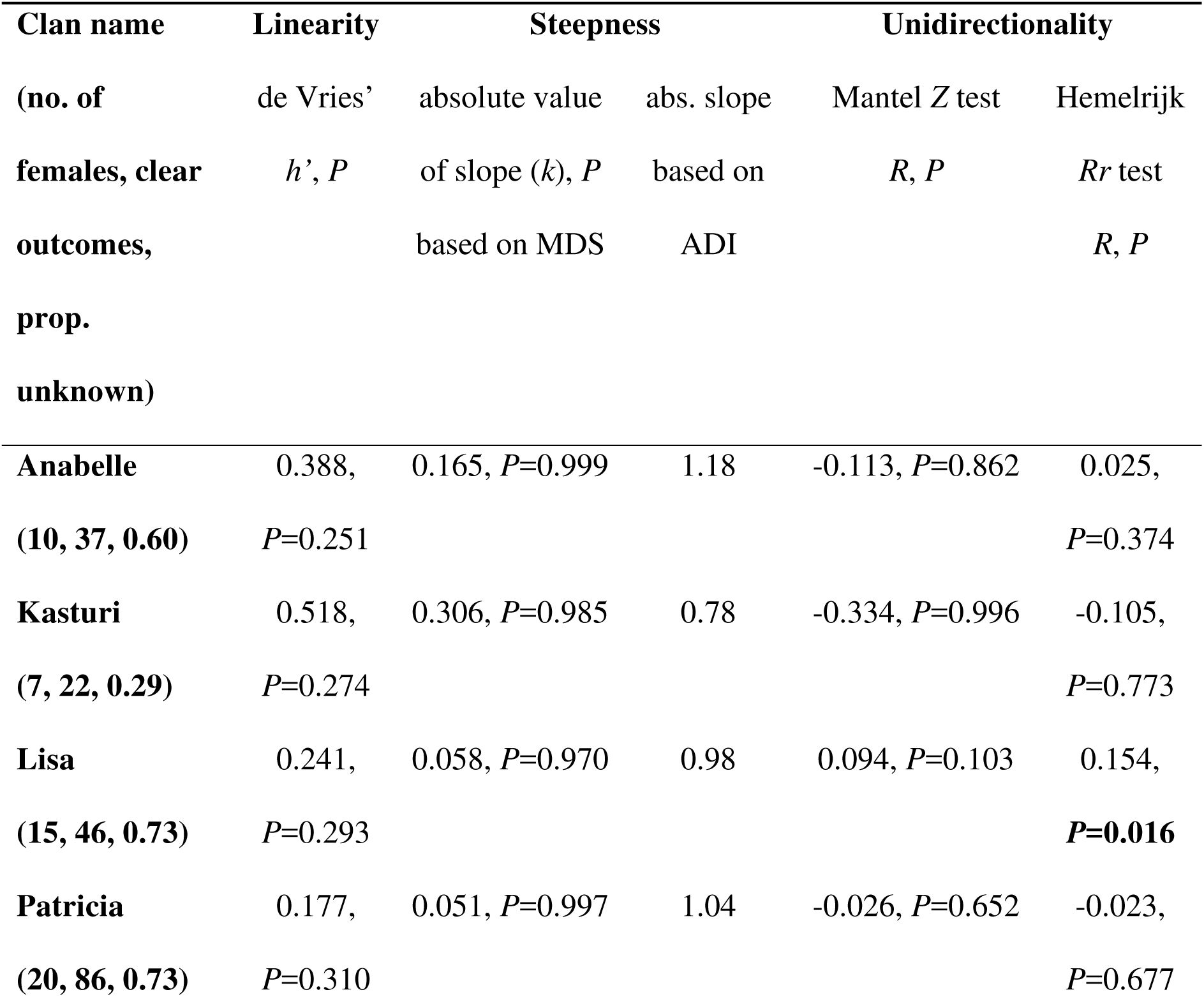
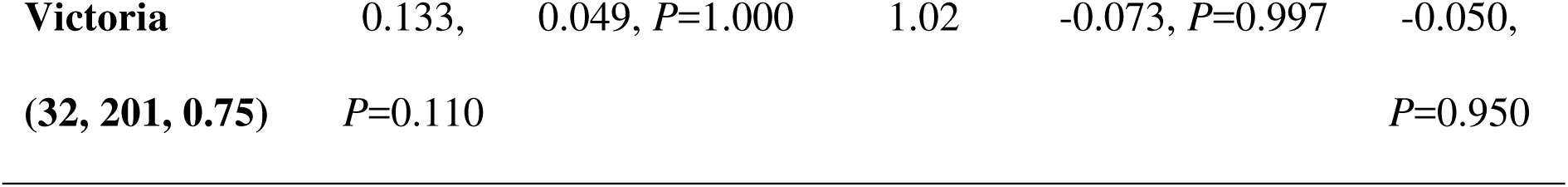
Quantification of within-clan dominance structure using measures of linearity and steepness of dominance hierarchy, and unidirectionality of dominance relationships in five common clans. Linearity of hierarchy was assessed using de Vries’ modified version of Landau’s *h* (*h*’, de Vries 1995) using 5000 permutations in SOCPROG. Steepness of hierarchy was assessed by testing the correlation between normalized modified David’s score (de Vries *et al*. 2006) and rank based on this dominance score. Steepness based on normalised average dominance index (ADI) is also shown for comparison. Unidirectionality was assessed by checking the correlation between the dominance matrix and its transposed matrix, using Mantel *Z* test of absolute reciprocity and Hemelrijk *Rr* test of relative reciprocity (correlation values and *P* are shown). Prop. unknown is the proportion of unknown relationships or empty cells in the dominance matrix.

**Table S4.** Results from analysis of triangle transitivity and triad census to study the orderliness of dominance within clans in five focal clans. P(*t*) is the proportion of transitive triads (out of the transitive and cyclic triads), *P* shows the statistical significance of P(*t*) based on 1100 randomisations (significant values marked in bold), and *t*tri is the triangle transitivity, which ranges up to 1 (maximum orderliness). The last three columns show the numbers of three triad motifs (double-dominant, double-subordinate, and pass-along; see Figure S2) and their statistical significance (probability of the observed value being higher than random in the case of double-dominant and double-subordinate, and that of the observed value being less than random in the case of pass-along) based on 1100 randomisations.

| Clan name | No. of triangles | $P(t)$ | $P$ | $t_{\text{tri}}$ | Double-dominant<br>obs. value ($P$) | Double-subordinate<br>obs. value ($P$) | Pass-along<br>obs. value ($P$) |
| --- | --- | --- | --- | --- | --- | --- | --- |
| **Anabelle** | 13 | 0.962 | 0.299 | 0.846 | 1 (0.994) | 11 (0.115) | 6 (0.069) |
| **Kasturi** | 12 | 0.875 | 0.226 | 0.5 | 5 (0.225) | 3 (0.681) | 5 (0.287) |
| **Lisa** | 11 | 0.955 | 0.265 | 0.818 | 25 (**0.037**) | 13 (0.575) | 16 (0.092) |
| **Patricia** | 31 | 0.855 | 0.161 | 0.419 | 39 (0.618) | 47 (0.309) | 53 (**0.043**) |
| **Victoria** | 142 | 0.931 | 0.000 | 0.725 | 240 (**0.002**) | 139 (0.826) | 247(**0.042**) |

**Table S5.** Summary of the outcomes of 475 independent within-clan dominance interactions. Description of dominance interactions Within-clan dominance

| Description of dominance interactions | Within-clan dominance |  |
| --- | --- | --- |
|  | No. of interactions | Percentage of all independent interactions |
| Interactions with clear winners | 465 | 97.9 |
| Interactions where there was no winner | 10 | 2.1 |
| Interactions where initiators won | 444 | 93.5 (95.5% of interactions with clear winners) |
| Interactions where recipients won | 21 | 4.4 (4.5% of interactions with clear winners) |
| Interactions where the initiator was older than the recipient | 356 | 74.9 (75.3% of interactions with clear winners) |
| Interactions where the initiator was younger than the recipient | 119 | 25.1 (24.7% of interactions with clear winners) |
| Interactions where the winner was older than the loser | 355 | 74.7 (76.3% of interactions with clear winners) |
| Interactions where the winner was younger than the loser | 110 | 23.2 (23.7% of interactions with clear winners) |

**Table S6.** Percentage of reversals against age (i.e., the percentage of decided interactions in which younger females won against older females), the total number of interactions with clear winners (*N*), and the absolute age differences between winners and losers when the former were older and when the former were younger. Results of Mann-Whitney *U* tests of the difference in these age differences in the two cases are shown. Significant *P* values are marked in bold. Values are given based on all the interactions with clear outcomes and also based on only unique dyads (in parentheses).

| Clan name | Reversals |  | Winner older than |  | Winner younger |  |  |  |
| --- | --- | --- | --- | --- | --- | --- | --- | --- |
|  | against age |  | loser |  | than loser |  |  |  |
|  | Percent | *N* | Absolute age | *N*1 | Absolute age | *N*2 | *MWU* | *P* |
|  |  |  | difference |  | difference |  |  |  |
| | | | average ± SD | | average ± SD | | | |
|  |  |  | years |  | years |  |  |  |
| **Anabelle** | 35.1% | 37 | 20.6 ± 15.49 | 24 | 4.9 ± 7.00 | 13 | 27.5 | **0.003** |
| | | | (20.6 ± 16.41) | (14) | (8.0 ± 9.70) | (6) | (22.0) | (0.099) |
| **Kasturi** | 22.7% | 22 | 26.7 ± 13.93 | 17 | 10.7 ± 11.14 | 5 | 16.0 | **0.038** |
| | | | (22.1 ± 13.12) | (11) | (10.7 ± 11.14) | (5) | (13.0) | (0.100) |
| **Lisa** | 26.1% | 46 | 22.9 ± 12.59 | 34 | 9.9 ± 8.59 | 12 | 95.5 | **0.007** |
| | | | (21.7 ± 13.48) | (23) | (10.1 ± 9.13) | (8) | (46.0) | **(0.038)** |
| **Patricia** | 23.3% | 86 | 15.3 ± 7.81 | 66 | 6.9 ± 5.07 | 20 | 221.0 | **<0.001** |
| | | | (15.3 ± 9.09) | (38) | (7.7 ± 5.68) | (15) | (131.5) | **(0.002)** |
| **Victoria** | 22.9% | 201 | 20.1 ± 13.87 | 155 | 12.1 ± 11.49 | 46 | 2264.5 | **<0.001** |
| | | | (19.7 ± 14.52) | (98) | (13.7 ± 12.20) | (34) | (1236.0) | **(0.025)** |

**Table S7.** The relationship between age and within-clan dominance. The first part has results of regression of dominance score (normalised Modified David’s Score) on the age of individuals (*N* is the number of females involved in dominance) are shown. *Rp* is the Pearson’s correlation coefficient. Significant *P* values based on 1000 permutations are marked in bold. The column *kage rank* is the slope of the relationship between normalized MDS and age-based rank (not the absolute age), thus representing the steepness of the age-based order. For comparison, the column *kage rank ADI* is the slope of the relationship between normalised average dominance index (ADI) and age-based rank.

| Clan name | $N$ | $R_p$ | $P$ | $k_{age\ rank}$ | $k_{age\ rank\ ADI}$ |
| --- | --- | --- | --- | --- | --- |
| Anabelle | 10 | 0.611 | 0.050 | -0.090 | -0.531 |
| Kasturi | 7 | 0.772 | **0.037** | -0.207 | -0.495 |
| Lisa | 15 | 0.745 | **0.001** | -0.043 | -0.519 |
| Patricia | 20 | 0.648 | **0.001** | -0.036 | -0.724 |
| Victoria | 32 | 0.594 | **<0.001** | -0.030 | -0.645 |

**Table S8.** The relationship between age of clan members and centrality measures calculated from female association networks in the five common clans: a) closeness centrality, and b) betweenness centrality. All females in Kasturi’s clan had the same centrality measures and were, therefore, not analysed for regression of these measures on age. Pearson’s correlation coefficient (*R*p) and *P* values based on 1000 permutations are shown.

| Clan | a) Degree |  | b) Closeness centrality |  | c) Betweenness centrality |  |
| --- | --- | --- | --- | --- | --- | --- |
| | $R_p$ | $P$ | $R_p$ | $P$ | $R_p$ | $P$ |
| **Anabelle** | -0.251 | 0.458 | 0.251 | 0.458 | -0.251 | 0.458 |
| **Kasturi** | - | - | - | - | - | - |
| **Lisa** | -0.166 | 0.609 | 0.166 | 0.609 | -0.183 | 0.508 |
| **Patricia** | -0.296 | 0.200 | 0.316 | 0.167 | -0.251 | 0.297 |
| **Victoria** | -0.334 | 0.034 | 0.334 | 0.034 | -0.246 | 0.196 |

**Table S9.**
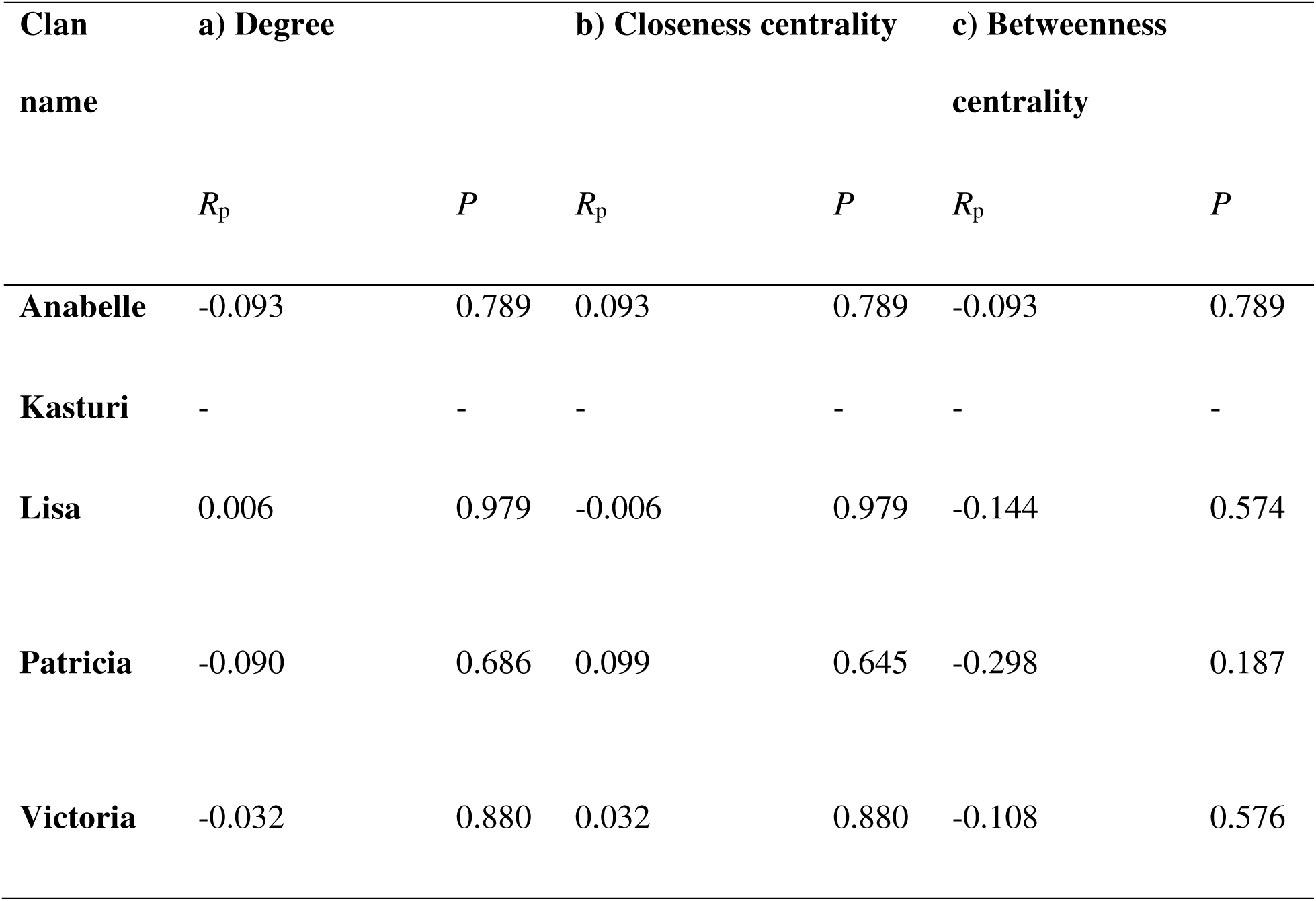
The relationship between dominance score (normalized MDS) of clan members and centrality measures calculated from female association networks in the five common clans: a) degree centrality b) closeness centrality, and c) betweenness centrality. All females in Kasturi’s clan had the same centrality measures and were, therefore, not analysed. Pearson’s correlation coefficient (*R*p) and *P* values based on 1000 permutations are shown.

| Clan<br>name | a) Degree |  | b) Closeness centrality |  | c) Betweenness<br>centrality |  |
| --- | --- | --- | --- | --- | --- | --- |
| | $R_p$ | $P$ | $R_p$ | $P$ | $R_p$ | $P$ |
| **Anabelle** | -0.093 | 0.789 | 0.093 | 0.789 | -0.093 | 0.789 |
| **Kasturi** | - | - | - | - | - | - |
| **Lisa** | 0.006 | 0.979 | -0.006 | 0.979 | -0.144 | 0.574 |
| **Patricia** | -0.090 | 0.686 | 0.099 | 0.645 | -0.298 | 0.187 |
| **Victoria** | -0.032 | 0.880 | 0.032 | 0.880 | -0.108 | 0.576 |

**Table S10.**
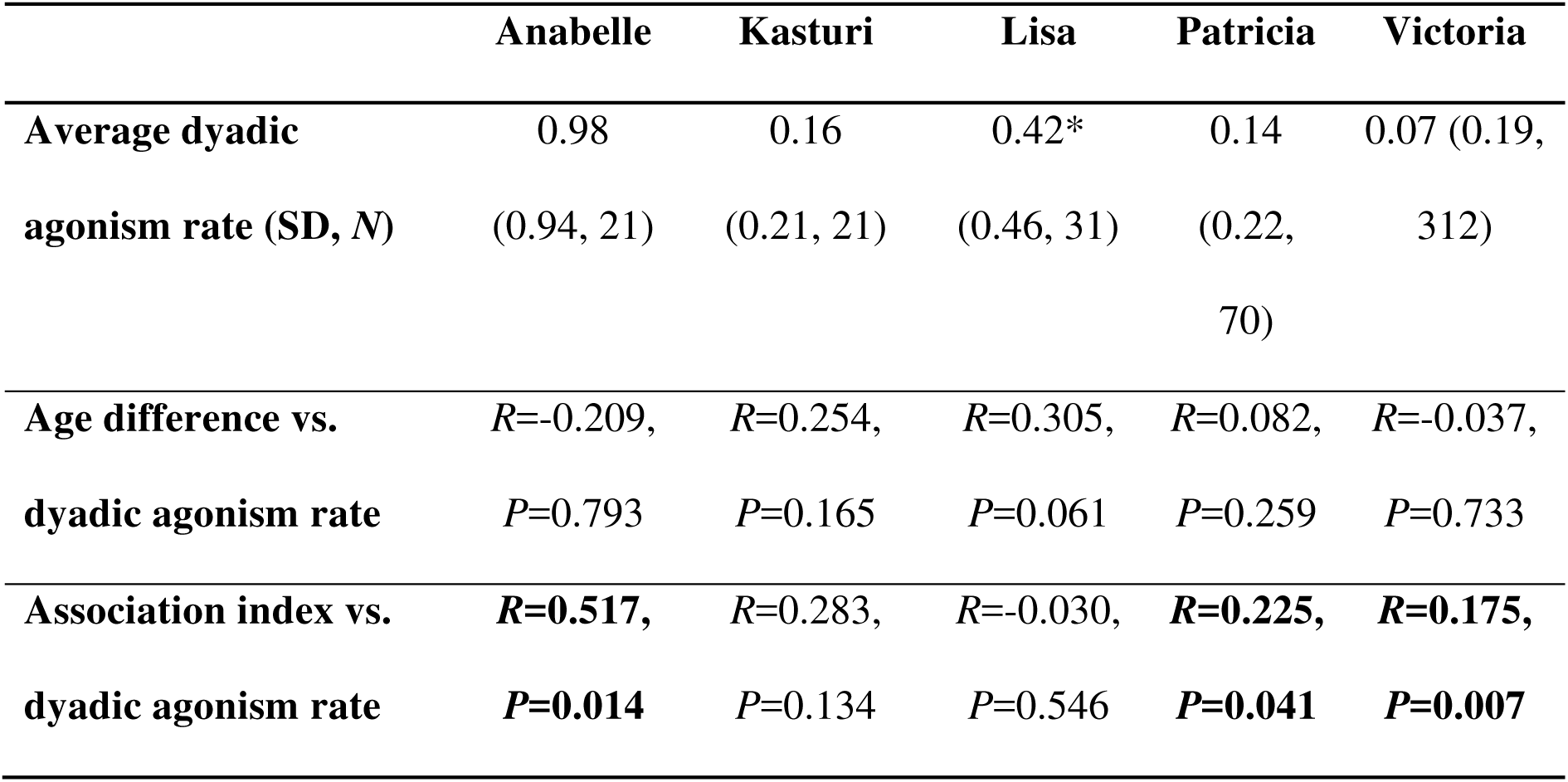
Dyadic rate of agonism (interactions/dyad/hour) within clans and the results of Mantel tests examining the relationship between dyadic rate of agonism and a) age difference, and b) social proximity (association index) in five common clans. The reported correlation coefficient is Pearson’s *R*. *Dyads seen for at least 55 minutes included in the calculation of the rate of agonism, unlike those seen for at least 60 minutes in other clans.

|  | **Anabelle** | **Kasturi** | **Lisa** | **Patricia** | **Victoria** |
| --- | --- | --- | --- | --- | --- |
| **Average dyadic** | 0.98 | 0.16 | 0.42* | 0.14 | 0.07 (0.19, |
| **agonism rate (SD, N)** | (0.94, 21) | (0.21, 21) | (0.46, 31) | (0.22, | 312) |
|  |  |  |  | 70) |  |
| **Age difference vs.** | *R*=-0.209, | *R*=0.254, | *R*=0.305, | *R*=0.082, | *R*=-0.037, |
| **dyadic agonism rate** | *P*=0.793 | *P*=0.165 | *P*=0.061 | *P*=0.259 | *P*=0.733 |
| **Association index vs.** | ***R*=0.517,** | *R*=0.283, | *R*=-0.030, | ***R*=0.225,** | ***R*=0.175,** |
| **dyadic agonism rate** | ***P*=0.014** | *P*=0.134 | *P*=0.546 | ***P*=0.041** | ***P*=0.007** |

## Part C: Supplementary Figures

**Figure S1.**
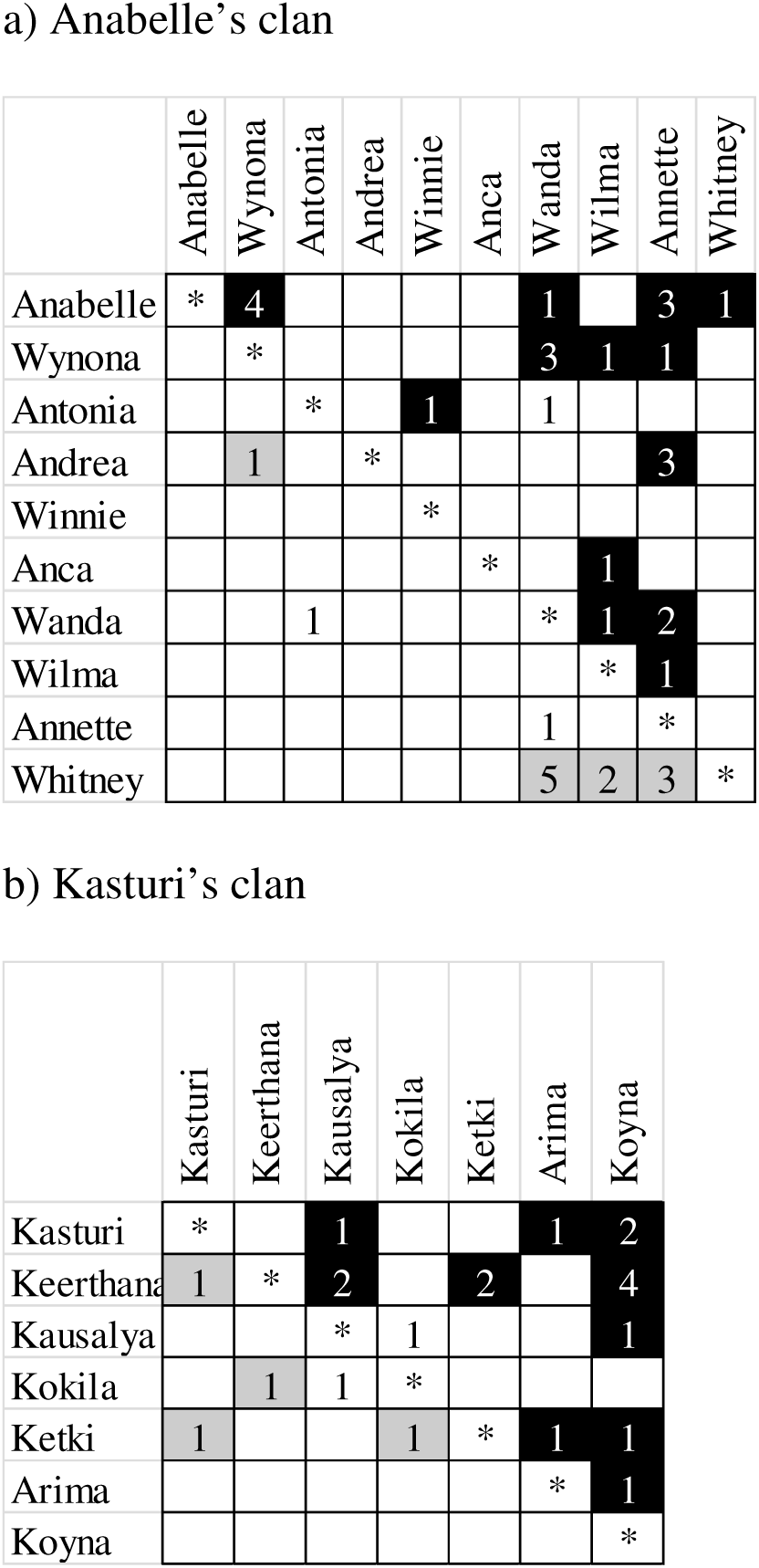
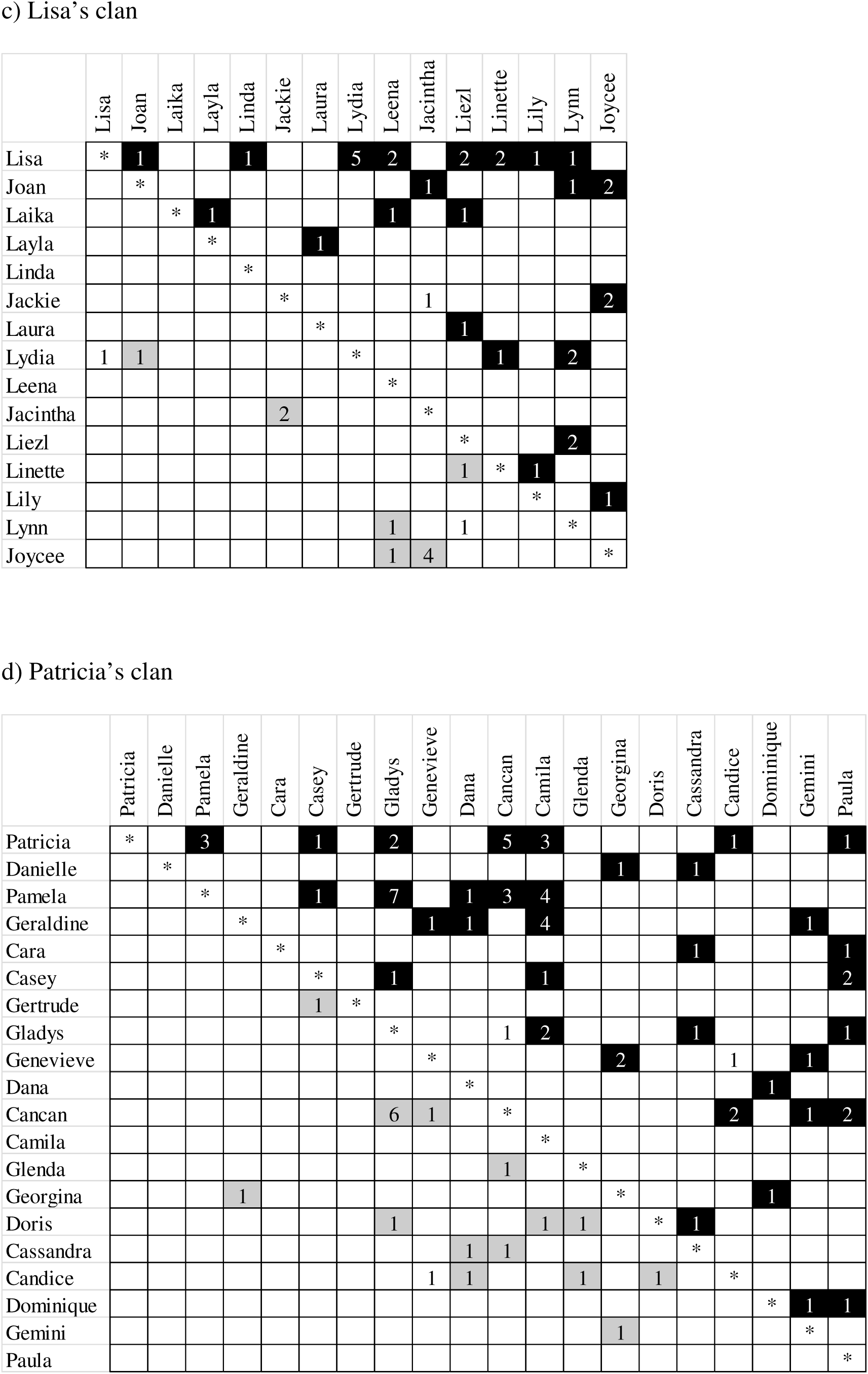
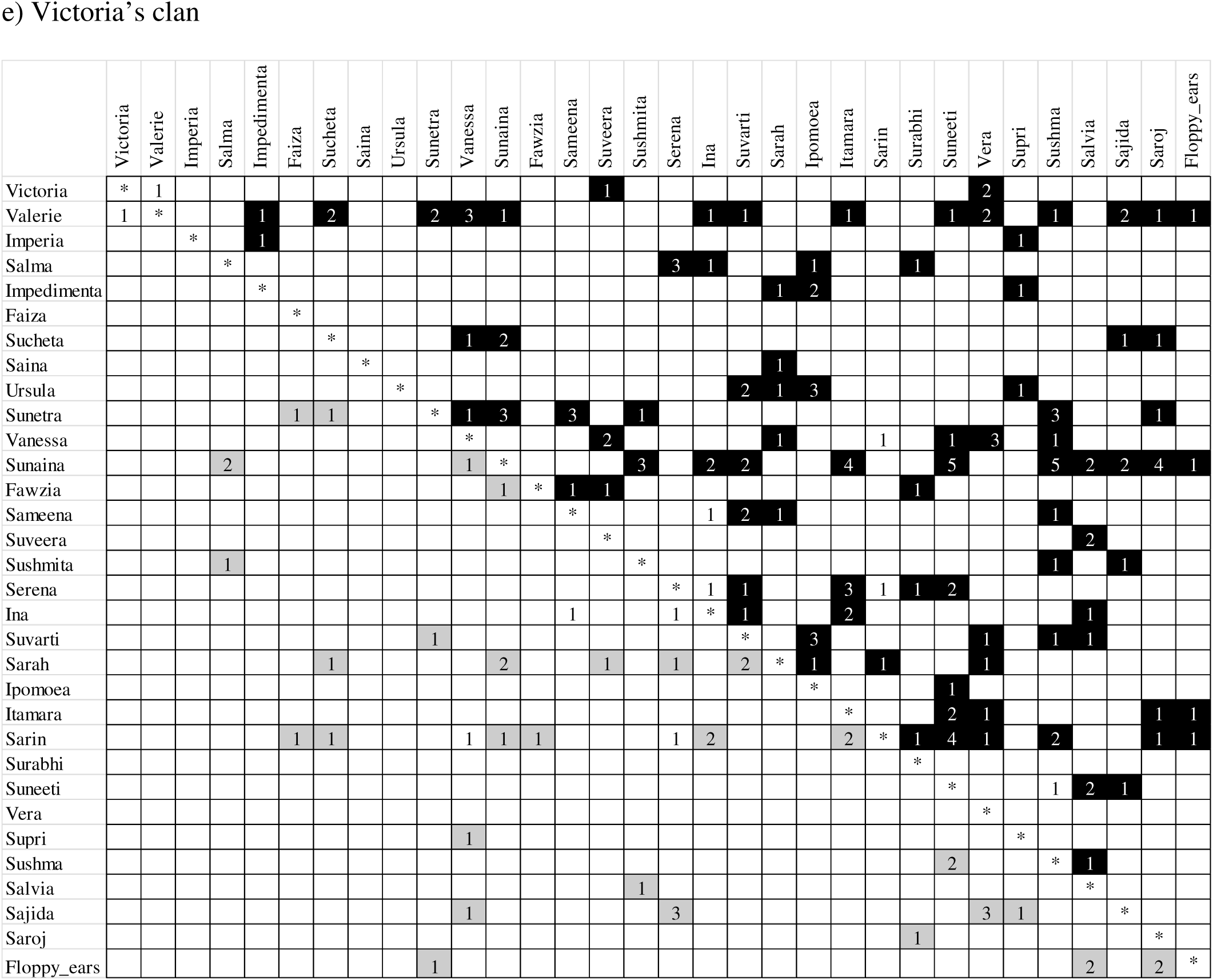
Dominance matrices showing numbers of all the independent agonistic interactions between adult females in the five focal clans. The rows represent the winners and the columns represent the losers of dominance interactions. Females are ordered by age, decreasing from left to right, and from top to bottom. Therefore, if dominance was based on age, one would expect to see most of the dominance in the upper right triangle. The black cells indicate instances when the older female won the majority of the agonistic interactions. Grey cells indicate instances when the younger female won the majority of the agonistic interactions.

**Figure S2.**
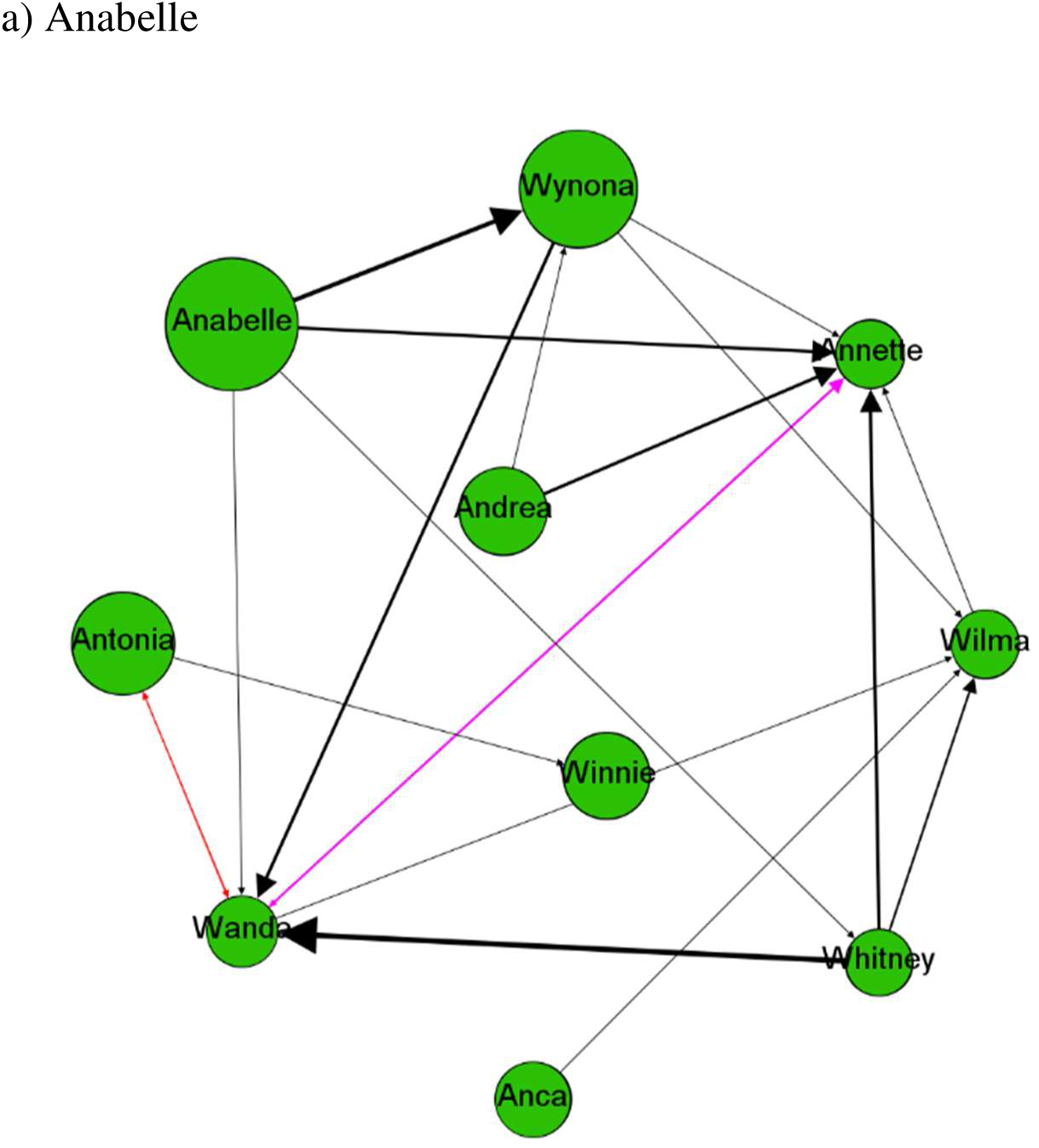
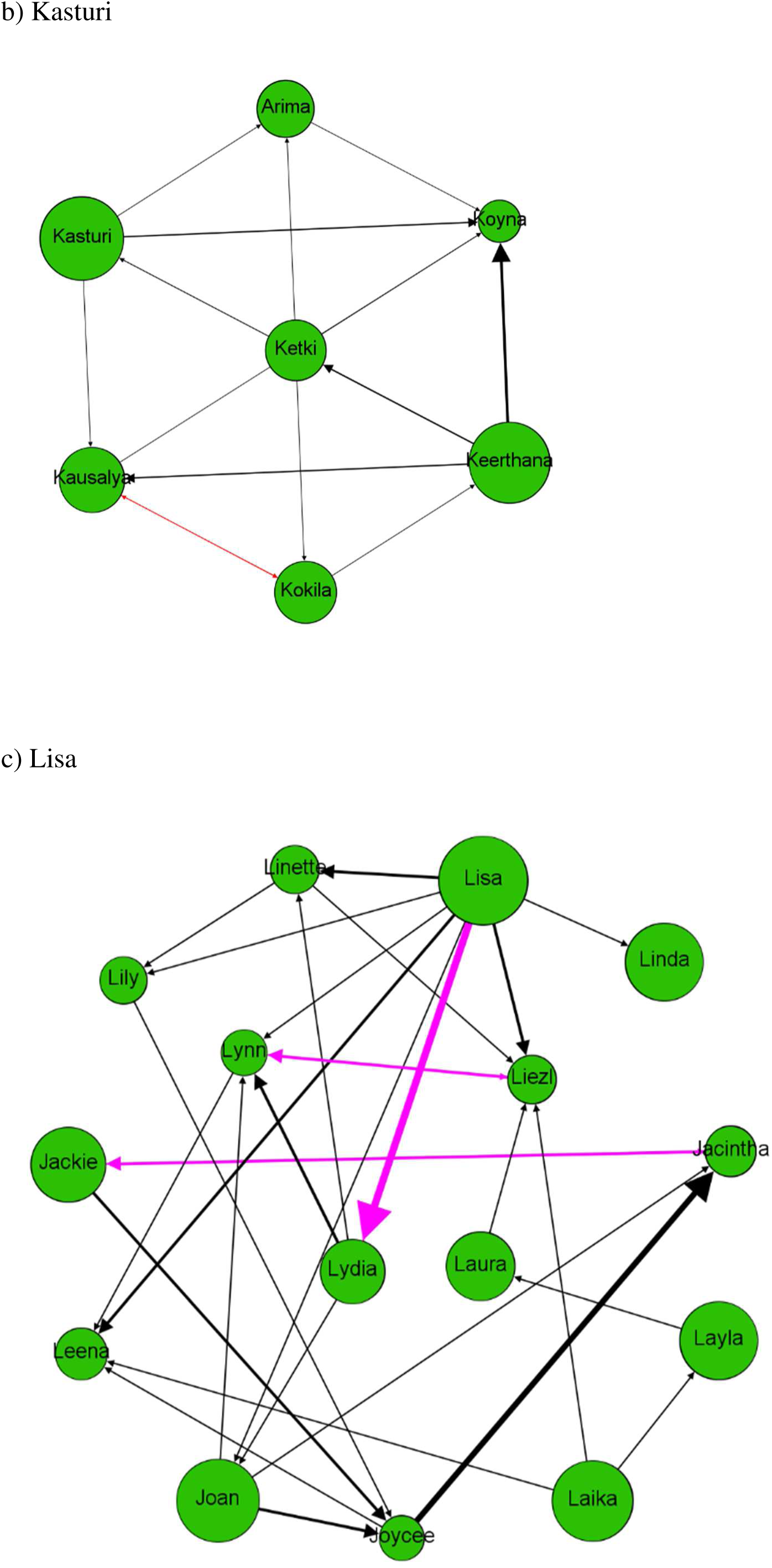
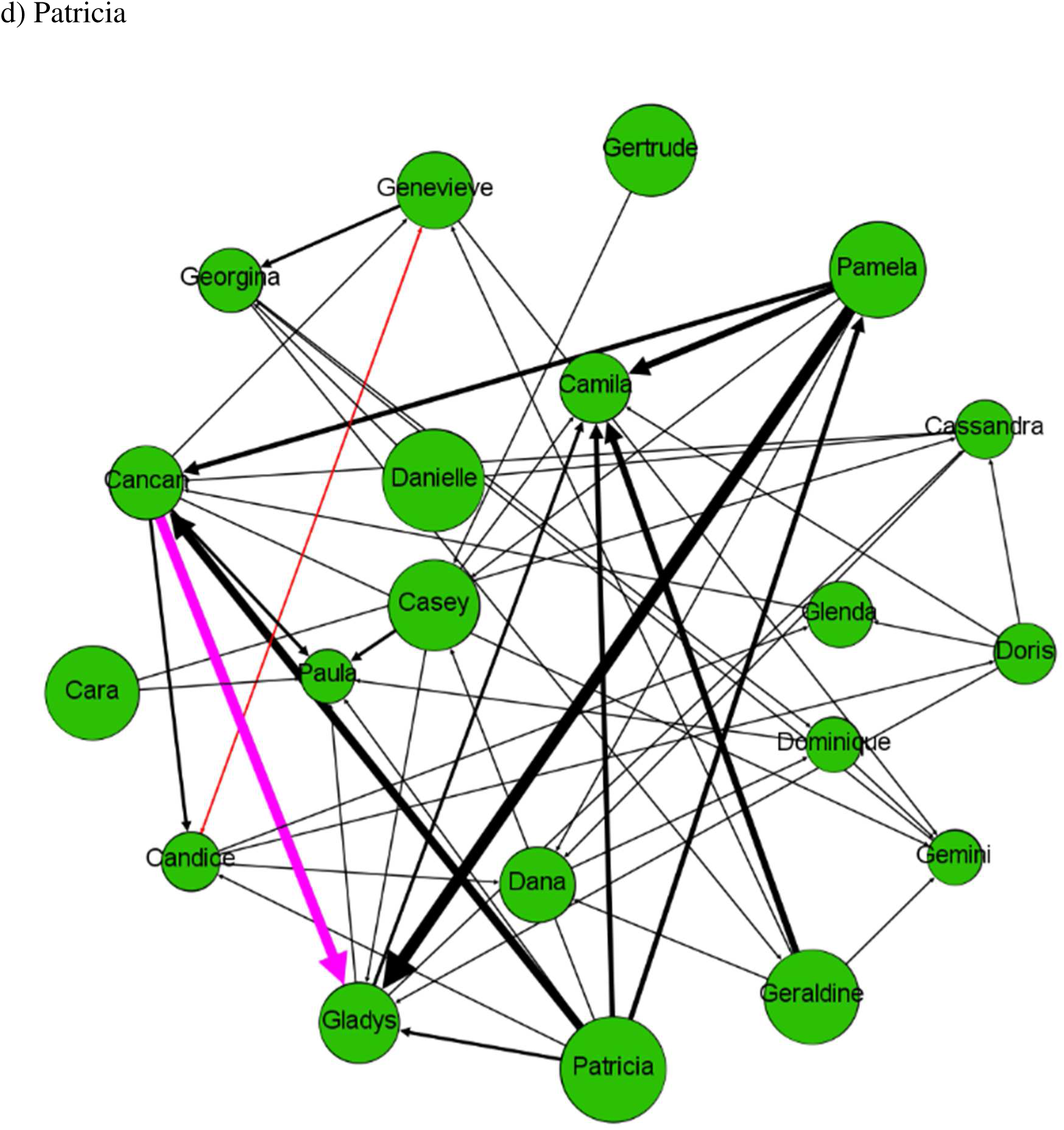
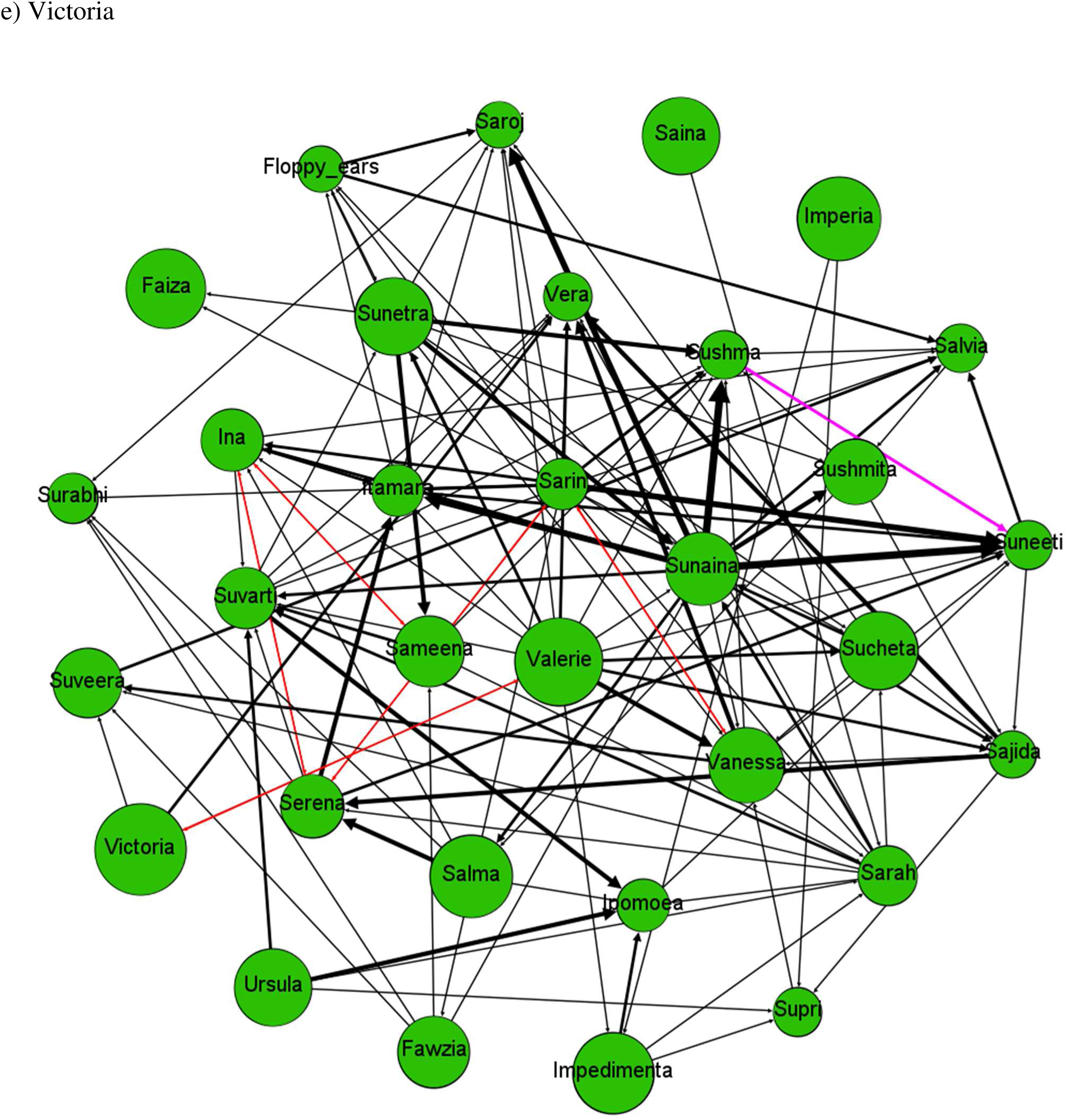
Within-group dominance networks (based on wins and losses) for the five common clans. Each node is a female and lines indicate agonistic interactions in the directions towards which the arrows point (for instance, an arrow from individual A to B indicates that A won against B). Nodes are sized within clans according to the age of females (larger nodes are older females) and thickness of the edges is proportionate to the number of interactions. Black lines are unidirectional relationships, where one individual won all the interactions, red lines indicate bidirectional relationships with equal wins, and pink lines indicate bidirectional relationships wherein one individual won the majority of the interactions (as indicated by the size of the arrow head).

